# Cortical circuit implementation of signed deviant detection with positive and negative prediction error neurons

**DOI:** 10.64898/2026.09.08.750236

**Authors:** Carlos A. Sanchez-Leon, Anand Suresh, Nicole H. Ershaghi, Carlos Portera-Cailliau

## Abstract

Predicting future events is fundamental for animals to adaptively interact with their environment, but whether and how the cerebral cortex processes predictive information remains an unresolved question. We combined in vivo two-photon calcium imaging in mouse somatosensory cortex with an oddball paradigm that decouples sensory stimulus properties from predictability context. We recorded neuronal responses in layer (L)2/3, bottom-up sensory inputs, and top-down prefrontal feedback. We find that L2/3 pyramidal neurons concurrently encode stimulus features and statistical context. We identify coexisting positive prediction error (PE) neurons that subtract predictions from incoming sensory inputs and negative PE neurons that do the opposite. This architecture represents a unified computational principle where error signals scale proportionally with the magnitude of mismatch across distinct stimulus features tested (amplitude, direction, omission). Furthermore, bottom-up pathways operate as unsigned novelty detectors, L2/3 circuit computes signed PE, and top-down projections signal a precision-weighted signed error, demonstrating hierarchical predictive processing.

## Introduction

The brain has been conceptualized as a predictive machine, constantly generating internal models of the world to anticipate and adapt to a changing environment (Clark, 2013). This predictive capacity relies on the computation of error signals, the discrepancies between internal expectations and actual inputs, which are then used by the brain to update its models. However, the circuit mechanisms that implement prediction in the cerebral cortex have not been fully characterized, and there is some debate as to whether true prediction even occurs at the circuit level (Carbajal and Malmierca, 2018; May, 2021).

Predictive processing is an influential theoretical framework (Rao and Ballard, 1999; Friston, 2005) that proposes the existence of a canonical cortical microcircuit (Douglas and Martin, 2004) implementing a common computation whereby bottom-up sensory information is continuously compared against top-down, intrinsically generated expectations (Bastos et al., 2012; Keller and Mrsic-Flogel, 2018). When a discrepancy occurs, the circuit generates a prediction error signal, which is then passed along the cortical hierarchy to update and refine the brain’s internal model, ultimately shaping adaptive behavior.

Predictive processing theories (Aizenbud et al., 2026) offer precise, experimentally testable hypotheses regarding the implementation of this essential cortical computation (Keller and Mrsic-Flogel, 2018): 1) Neuronal responses to sensory inputs are not merely a passive representation of bottom-up stimulus features (e.g., stimulus-specific adaptation, SSA) (Adibi and Lampl, 2021) but are instead actively modulated by internal expectations (de Lange et al., 2018). 2) The cortex must possess two different computational units: prediction neurons, primarily located in deep cortical layers and sending feedback projections to lower-order areas (Bastos et al., 2020), and prediction-error (PE) neurons, primarily located in superficial layers that send feed-forward projections to higher-order areas (Hamm et al., 2021; Furutachi et al., 2023). In essence, PE neurons compute the difference between bottom-up inputs and top-down predictions and send any discrepancy (i.e., the error) up the hierarchy to update the brain’s internal model. 3) Because pyramidal neurons fire sparsely (Crochet et al., 2011; Wang et al., 2022), a single error-encoding population is unlikely to bidirectionally signal the direction of the deviation, i.e., whether a sensory input is larger or smaller than expected. Thus, there must exist two distinct categories of PE neurons: positive PE (pPE) neurons, which increase their activity when a sensory stimulus is larger than expected, and negative PE (nPE) neurons, which increase their activity when the stimulus is smaller than expected or completely absent (Fiser et al., 2016; Lao-Rodríguez et al., 2023; Leonardon et al., 2025; Yaron et al., 2025). 4) The magnitude of the PE signal is weighted by the confidence of the internal model (Friston, 2018; Hodson et al., 2024). 5) the PE computation is distributed across the cortical hierarchy and shaped by local interneurons (Attinger et al., 2017; Parras et al., 2017; Casado-Román et al., 2020; Ding et al., 2026; Tsukano et al., 2026).

Over the past several years, empirical evidence has supported some, but not all, of these claims. In particular, the existence of pPE and nPE neurons has not been unequivocally demonstrated, and a fully elaborated canonical microcircuit for prediction has not yet emerged. Moreover, whether the cerebral cortex even implements predictive coding remains a subject of intense debate (Walsh et al., 2020; Hodson et al., 2024). A limitation of previous empirical studies of prediction is that the above-named hypotheses have been considered in isolation and have been tested across different experimental paradigms, animal models, and recording techniques (Parras et al., 2017; Casado-Román et al., 2020; Hamm et al., 2021; Garner and Keller, 2022; Heilbron et al., 2022; Furutachi et al., 2023; Ding et al., 2026).

To overcome these limitations, we implemented an oddball paradigm designed to systematically isolate a single computational variable at a time, either the sensory input or the prediction. We then used two-photon calcium imaging in mouse vibrissal somatosensory cortex (vS1) to record the responses of individual neurons to the exact same sensory input across different predictability contexts (unexpected, expected, or predictively neutral), as well as within an identical predictive context across varying sensory inputs (larger or smaller than expected). By isolating statistical context from sensory variability, we provide direct evidence that vS1 simultaneously encodes whisker deflection amplitude and its statistical context through dedicated PE neurons. We reliably observed predictive ensembles, consisting of pPE and nPE neurons whose complementary activity was governed by a mirrored connectivity motif. These ensembles perform a fundamental computation that generalizes across stimulus features to also process complete sensory omissions, as well as changes in whisker direction. Lastly, our findings reveal a hierarchical transformation from unsigned deviant detection in bottom-up L4 and posteromedial thalamic nucleus (POm) projections, to signed PEs in L2/3, and ultimately a precision-weighted, signed PE feedback from secondary motor cortex (M2).

## RESULTS

In this study, we set out to systematically test the existence of the basic elements of a prediction circuit (Keller and Mrsic-Flogel, 2018). Such a circuit should contain a bottom-up sensory signal, a L2/3 circuit composed of both pPE and nPE neurons, and a top-down prediction signal (**Fig. 1a**).

**Fig. 1.**
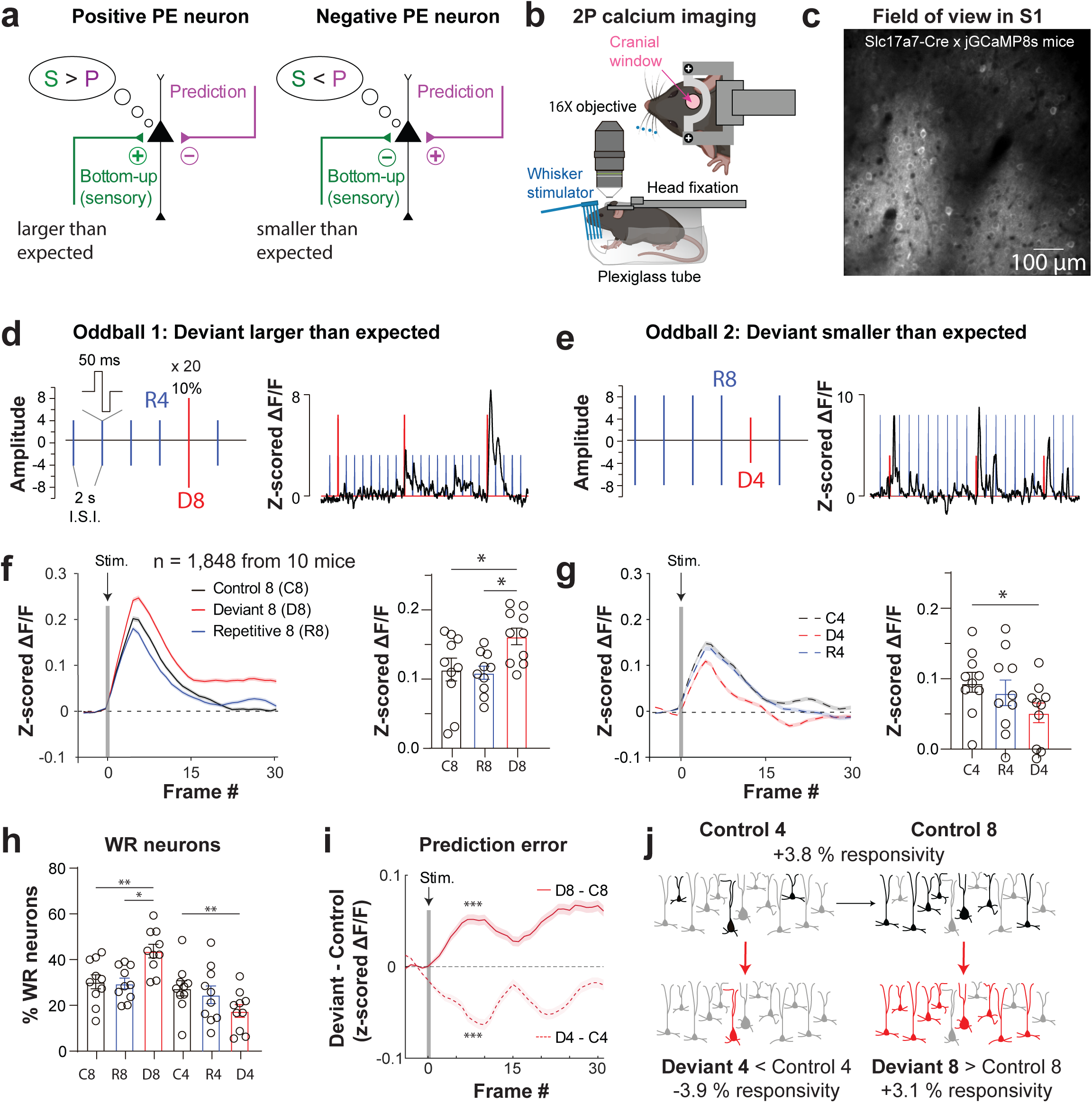
L2/3 pyramidal neurons simultaneously encode whisker amplitude and predictability context. **a)** Schematic of connectivity motifs for positive and negative prediction error (PE) neurons. Putative excitatory and inhibitory connections are represented with + and – symbols, respectively. **b)** Cartoon of experimental setup for head-fixed 2-photon (2P) calcium imaging during whisker stimulation. **c)** Representative *in vivo* two-photon calcium imaging field of view showing L2/3 pyramidal neurons in a transgenic Slc17a7-Cre x jGCaMP8s mouse. **d)** Schematic of the oddball protocol #1 where the Deviant is larger than expected (*left*), and representative z-scored ΔF/F calcium trace from individual neurons (*right*). **e)** Same as *d*, but for oddball protocol #2, where Deviant is smaller-than expected. **f)** Mean evoked population activity (z-scored ΔF/F) of neurons that were WR relative to Amplitude 4 and 8 stimuli (n= 1,848 neurons from 10 mice) (*left*) and corresponding statistical comparison per mouse (*right*) for Protocol #1 (Deviant = Amplitude 8) across three predictability contexts: Control (C, black), Repetitive (R, blue), and Deviant (D, red). **g)** Same as *f* but for Protocol #2 (Deviant = Amplitude 4). (Two-way ANOVA, Context p= 0.326, Amplitude p= 0.004, Context x Amplitude interaction p= 0.009, N= 10 mice). **h)** Percentage of recruited WR neurons per mouse across different contexts (Two-way ANOVA, Context p= 0.354, Amplitude p= 0.006, Context x Amplitude interaction p= 0.002, N= 10 mice). **i)** Subtracted mean evoked population activity isolating the prediction error (PE) modulation (mean evoked response to Deviant minus its corresponding Control; D - C) for greater-than-expected (D8 – C8, solid line) and smaller-than-expected violations (D4 – C4, dashed line) (Linear Mixed-Effects Model, Context p<0.001, Amplitude p<0.001, Context x Amplitude interaction p<0.001, n= 1,848 neurons from 10 mice). **j)** Cartoon of the dual modulation of cortical ensembles by amplitude and predictability context Left → Right: Larger amplitudes recruit more neurons. Top → Bottom: Larger/smaller deviants recruit more/fewer neurons, respectively. *p<0.05; **p<0.01; ***p<0.001. Shaded areas and error bars represent SEM.

### L2/3 pyramidal neurons encode whisker amplitude

We first characterized how L2/3 pyramidal neurons in vS1 (barrel cortex) process stimuli of different amplitudes in a predictively neutral context. The C2 barrel was first localized with intrinsic signal imaging (**Extended Data Fig. 1a**) and then we performed two-photon calcium imaging in awake, head-fixed transgenic mice expressing GCaMP8s in excitatory neurons (Slc17a7-jGCaMP8s; N= 10 mice; **Fig. 1b-c**). We recorded responses of L2/3 neurons to brief (50 ms) whisker deflections at different randomized amplitudes (0.18, 0.30, 0.53, 0.68, or 0.73 mm) generated by delivering different intensities (2, 4, 6, 8, or 10 V) to a piezoelectric actuator (**Extended Data Fig. 1b-c**). Whisker deflections were delivered every 2 s, a prolonged interval that minimized SSA (Adibi et al., 2013; Musall et al., 2017). We quantified mean response amplitude, responsivity, and maximum response amplitude (**Extended Data Fig. 1d-f**; see Methods). At a population level, whisker-responsive neurons (WR; n= 1,578 neurons) showed increased mean activity to deflections of larger amplitudes (**Extended Data Fig. 1d**). We observed a significant correlation of stimulus amplitude with mean evoked activity (**Extended Data Fig. 1g**), as well as with the percentage of recruited WR neurons (**Extended Data Fig. 1h**). Because L2/3 neurons in vS1 typically exhibit low firing rates and respond with low probability to whisker stimuli (Brecht et al., 2003; Kerr et al., 2007; Crochet et al., 2011), we examined whether the increase in mean evoked activity was due to an increase in the number of trials that the neurons responded to (i.e., a change in stimulus responsivity), an increase in maximum evoked response, or both. We found a significant correlation between responsivity (R^2^= 0.92; p= 0.01), but not with maximum response (R^2^= 0.39; p= 0.26), suggesting that stimulus amplitude is encoded by the recruitment probability rather than the magnitude of the neuronal response (**Extended Data Fig. 1i-j**).

Furthermore, neurons recruited by a particular amplitude showed smaller responses to other amplitudes (including larger ones), resulting in sharp tuning curves and a skewed selectivity index (SI) distribution (**Extended Data Fig. 1k-l**; mean SI= 0.60). Consistent with this high degree of specificity, only a small fraction of neurons exhibited responses across multiple amplitudes (**Extended Data Fig. 1m**). Finally, Support Vector Machine (SVM) classifiers trained on neuronal activity showed similar high decoding accuracy for all whisker deflection amplitudes (**Extended Data Fig. 1n**). Interestingly, when the decoder misclassified an amplitude, it predominantly categorized it as the most similar adjacent amplitudes (**Extended Data Fig. 1o**). These findings demonstrate that L2/3 pyramidal neurons encode stimulus amplitude as a discrete variable recruiting largely segregated functional ensembles while maintaining a structured sensory representation where closer amplitudes share more similarity.

### L2/3 pyramidal neurons encode predictability context

To determine whether L2/3 pyramidal neurons encode internal expectations about sensory inputs, we next recorded their responses to the same stimulus amplitude across distinct predictability contexts: 1) an ‘Oddball 1’ block in which Amplitude 4 was the repetitive stimulus (R4) and Amplitude 8 was the Deviant stimulus (D8) at 10% probability (D8 in R4 context; **Fig. 1d**); 2) a flipped ‘Oddball 2’ block (D4 in R8 context; **Fig. 1e**); 3) a Control (C) block (as in **Extended Data Fig. 1c**) in which different amplitudes appeared in random order with equal probability (20% each). This protocol allowed us to compare responses to the exact same physical stimulus under three levels of expectation: highly predictable (R4 in Oddball 1), unexpected violation (D4 in Oddball 2), and contextually neutral (C4 in Control block).

We focused our analysis on L2/3 neurons that responded to any stimuli of Amplitude 4 or 8 in any context. We found that individual L2/3 neurons exhibited robust responses to Deviants regardless of the direction of the amplitude change (**Fig. 1d-e, traces**). At a population level, we observed significantly larger responses to D8 (red) compared to their R8 (blue) or C8 (black) counterparts (**Fig. 1f and Extended Data Fig. 2a**). Conversely, the responses to D4 were significantly smaller than their corresponding R4 and C4 responses (**Fig. 1g and Extended Data Fig. 2b**). Accordingly, D8 and D4 stimuli recruited a greater and smaller percentage of WR neurons, respectively (**Fig. 1h**), and the change in their mean evoked activity was primarily driven by a change in responsivity rather than in maximum response (**Extended Data Fig. 2c-d**). By subtracting the C response from the D response, we isolated the true PE signal (Parras et al., 2017; Casado-Román et al., 2020) (**Fig. 1i; Extended Data Fig. 3**). Note that C and R neuronal responses were similar in our recordings, both as far as magnitude and neuronal recruitment, suggesting that the long inter stimulus interval (2s) minimizes potential repetition effects (Musall et al., 2017; Parras et al., 2017).

To determine if the vS1 population carries sufficient information to discriminate the same stimulus in the three different contexts, we trained multiple SVM classifiers. We found high decoding accuracy (>70%) for all contexts across both Amplitude 4 and 8 comparisons (**Extended Data Fig. 2e**), indicating that sensory representations in vS1 are dynamically reorganized as a function of statistical predictability. Furthermore, to control for potential probability-dependent effects (20% vs 10% probability in Control vs. Deviant contexts), we performed additional recordings using an expanded Control block consisting of 10 randomized amplitudes. We observed similar functional dynamics of evoked responses in L2/3 neurons (n= 2,379 neurons, 8 mice; **Extended Data Fig. 4a-c**), which once again showed high selectivity for their preferred amplitudes (**Extended Data Fig. 4d-e)**. Crucially, the direction of the PE responses to Deviants remained consistent: a significantly larger response to D8 than to 10%-occurrence C8 and R8, and a smaller response to D4 than to C4 and R4 (**Extended Data Fig. 4f-g**).

Thus, unexpected whisker inputs are strongly modulated bidirectionally, differentially signaling larger- vs. smaller-than-expected amplitudes, even after controlling for repetition-induced effects, and probability of stimulus occurrence. We conclude that L2/3 pyramidal neurons simultaneously encode stimulus amplitude and predictability, such that vS1 does not merely represent stimulus amplitude but can bidirectionally signal deviations from expected amplitudes (**Fig. 1j**).

### L2/3 pyramidal neurons compute signed prediction errors (PE) via positive (pPE) and negative (nPE) functional ensembles

To determine if this deviant detection/prediction capability depends on specialized neuronal ensembles, we identified neurons that were strongly modulated by the Deviant stimulus (see Methods). Neurons selective for D8 (17.8% of WR neurons) showed a significant increase in evoked activity compared to R/C conditions (**Fig. 2a**, solid line; **Extended Data Fig. 5a-b**), while those selective for D4 (15.5% of WR neurons) exhibited a decrease (**Fig. 2b**, dashed line; **Extended Data Fig. 5c-d**). Interestingly, despite being selected for their response to a particular Deviant amplitude (either D8 or D4), these same neurons were inversely modulated by the opposite Deviant, although to a lesser degree (**Fig. 2a**, dashed line; **Fig. 2b**, solid line; **Extended Data Fig. 5a-d**), effectively acting as bidirectional PE neurons. As in our previous population-level analysis, the mean evoked activity changes were mainly driven by a change in responsivity across trials (**Extended Data Fig. 5e-f**).

**Fig. 2.**
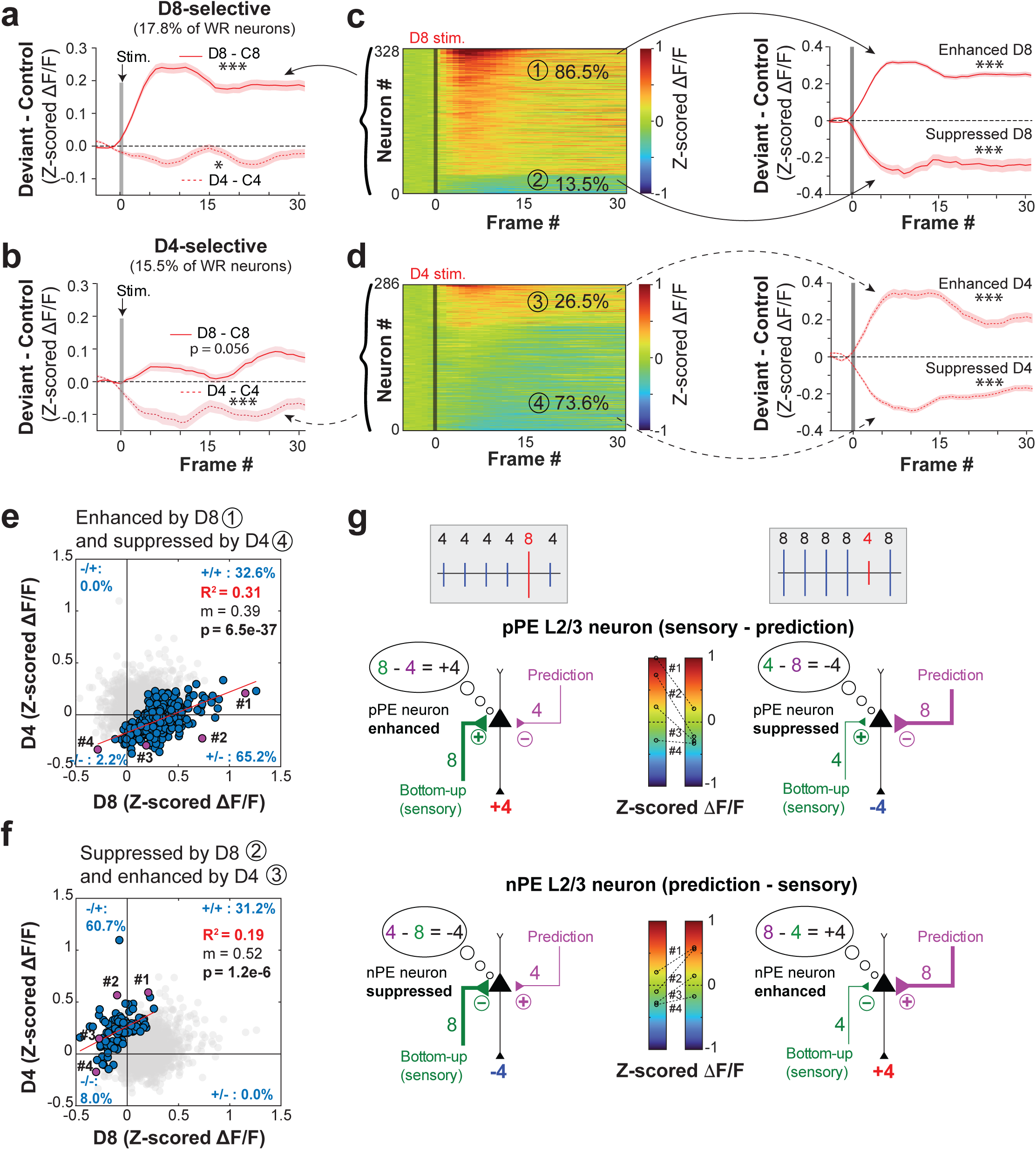
L2/3 pyramidal neurons compute signed prediction errors (PE) via positive (pPE) and negative (nPE) functional ensembles. **a)** Subtracted mean evoked PE activity (D – C, z-scored ΔF/F) of D8-selective neurons to D8 (solid line) or to D4 (dashed line) (Linear Mixed-Effects Model, Context p<0.001, Amplitude p<0.001, Context x Amplitude interaction p<0.001, n= 328 neurons from 10 mice). **b)** Same as *a* but for D4-selective neurons (Linear Mixed-Effects Model, Context p<0.003, Amplitude p<0.001, Context x Amplitude interaction p<0.001, n= 286 neurons from 10 mice). **c)** Left, single-neuron mean evoked activity (z-scored ΔF/F) of D8-selective neurons in response to D8, sorted by max value. Right, mean evoked PE activity (D8 – C8) of enhanced ① and suppressed ② subpopulations (Linear Mixed-Effects Model, enhanced: Context p<0.001, Amplitude p<0.001, Context x Amplitude interaction p<0.001, n= 285 neurons from 10 mice. Suppressed: Context p<0.001, Amplitude p<0.001, Context x Amplitude interaction p<0.001, n= 43 neurons from 10 mice). **d)** Same as *c* but for D4-selective neurons (Linear Mixed-Effects Model, enhanced ③: Context p<0.001, Amplitude p<0.001, Context x Amplitude interaction p<0.001, n= 76 neurons. Suppressed ④: Context p<0.001, Amplitude p<0.001, Context x Amplitude interaction p<0.001, n= 210 neurons). **e)** Cross-deviant correlation scatter plot for the dominant/majority ensembles (enhanced activity by D8 and suppressed by D4). Percentages indicate the relative distribution of neurons across quadrants. Example neurons in purple are shown in g. **f)** Same as *e* but for the minority ensembles (suppressed by D8 and enhanced by D4). **g)** Schematic illustrating the proposed arithmetic calculation made by pPE and nPE neurons. Mirroring connectivity motifs subtract either top-down predictions from incoming sensory inputs (pPE neurons) or incoming sensory inputs from top-down predictions (nPE neurons), determining the sign of the error and the recruitment of specialized ensembles across oddball contexts. Heatmap of mean evoked activity from example neurons in panels *e* and *f.* *p<0.05; ***p<0.001. Shaded areas in *a-d* represent SEM.

To validate the necessity and sufficiency of this subpopulation, we trained SVM classifiers exclusively on the responses of these deviant-selective PE neurons and found high (>70%) overall decoding accuracy for all contexts (**Extended Data Fig. 5g**, red bar), exceeding that of an equal number of randomly selected neurons (**Extended Data Fig. 5g**, light red bar). Removal of the PE neurons (a relatively small fraction of the entire population) significantly reduced context decoding accuracy (grey bar), while removing an equal number of random neurons did not (white bar), reaching similar accuracy levels as the entire population (**Extended Data Fig. 5g**, dark grey bar). As further evidence that our selection criteria accurately identified the core PE neurons driving context discrimination, we identified the subpopulation of neurons with the highest weights (top 5%) in the decoder trained on the entire population. While these neurons maximize separability among the three contexts without a priori bias toward any single condition, their responses mirrored those of our selected PE ensembles (**Extended Data Fig. 5h**): increased evoked activity for D8 and decreased for D4 relative to the other conditions. The high degree of overlap between these populations (**Extended Data Fig. 5h**) likely explains why eliminating the PE subpopulation from the decoder led to such a dramatic decrease in accuracy, and validates our subsequent analysis on these specific ensembles.

While the average activity was bidirectionally modulated, individual neural responses revealed a functional heterogeneity within these PE ensembles (**Fig. 2c-d**, left heatmaps; see Methods): Deviant stimuli enhanced the activity of some neurons but suppressed it for others. ‘Enhanced’ neurons showed significantly higher activity for both D8 and D4 (groups ①and ③), while ‘suppressed’ neurons showed a reduction in activity (groups ② and ④; **Fig. 2c-d**, right traces). Furthermore, the magnitude of net activity modulation by the Deviant was comparable across both enhanced and suppressed populations, irrespective of the specific stimulus amplitude, indicating that the overall increase observed for D8-selective PE neurons (**Fig. 2a**, solid line) is driven by a higher proportion of ‘enhanced’ neurons (86.5 vs. 13.5%), whereas the decrease observed for D4-selective PE neurons (**Fig. 2b**, dashed line) reflects the dominant effect of ‘suppressed’ neurons (26.5 vs. 73.6%).

We next performed cross-deviant correlation analyses between the responses of individual neurons to D8 and D4 stimuli. We found that neurons in each PE ensemble (D8 selective or D4-selective) clustered into two well-differentiated functional groups (**Extended Data Fig. 5i**) and their responses across Deviants showed very low correlations (R^2^ = 0.01 and 0.08). Grouping neurons based on their modulation polarity (either always enhanced or always suppressed by both Deviants; **Extended Data Fig. 5j**, left and right, respectively) also showed low correlations (R^2^ = 0.14 and 0.02). In fact, the correlation was negative, indicating that neurons that increase their activity to a particular Deviant amplitude, decrease it to the other. This suggests that these functional PE ensembles bidirectionally signal the error rather than responding non-specifically to any deviant. Indeed, when we grouped neurons based on the dominant/majority ensemble from each deviant, i.e., enhanced activity for D8 and suppressed for D4, we observed a much stronger positive correlation (R^2^ = 0.31; **Fig. 2e**). Even when grouping PE neurons according to the minority ensemble, we observed a good positive correlation (R^2^ = 0.19; **Fig. 2f**). These functional profiles suggest that, despite their opposite contextual modulation (enhanced response to one deviant and suppressed to the other), PE neurons perform a common computation and/or have a shared input architecture.

These results align very well with previous formulations of predictive coding in the neocortex, in which the population computes a simple arithmetic calculation (Keller and Mrsic-Flogel, 2018). Under this scheme (**Fig. 2g**), the opposite modulation observed experimentally is a direct consequence of two mirroring connectivity motifs. Specifically, pPE neurons perform a Sensory -Prediction subtraction (**Fig. 2g**, top) and thus exhibit enhanced activity when sensory input exceeds prediction (i.e., D8 > R4; top left) but suppressed activity when sensory input falls below prediction (D4 < R8; top right). pPE neurons are therefore driven by an excitatory sensory input and an inhibitory prediction. Conversely, nPE neurons are driven by an inhibitory sensory input and an excitatory prediction, thus performing a Prediction - Sensory calculation (**Fig. 2g**, bottom), so they are suppressed when sensory input exceeds prediction (D8 > R4; bottom left) and enhanced when sensory input falls below prediction (D4 < R8, bottom right).

### Shared prediction but distinct sensory inputs define pPE and nPE functional ensembles

The above experiment provides strong evidence for the existence of pPE and nPE neurons in a scenario in which we compared neuronal responses to different sensory inputs (D8 or D4) during different prediction contexts (R4 and R8, respectively). If this model is correct, the same sensory input (D4) should be differentially modulated depending on whether the prediction is of a higher or lower expected amplitude, with proportional recruitment of enhanced and suppressed neurons. To test this, we implemented a variation of our oddball protocol where D4 is embedded in a predictable R2 context (and vice-versa, D2 embedded in a predictable R4 context; **Extended Data Fig. 6a**). We tracked the same neurons longitudinally during this new protocol, as well as during the original oddball paradigm (D4-R8 vs. D8-R4; **Extended Data Fig. 6b**, counterbalanced). This design allowed us to examine how the same sensory input (D4) was modulated by opposing predictive contexts of either smaller (R2) or larger (R8) amplitude. Importantly, it also allowed us to test the reciprocal computational axis: how these same neurons respond to distinct sensory deviations (D2 vs. D8) when the internal prediction remains constant (R4).

When a D4 stimulus followed a R2 sequence (a larger-than-expected violation), D4afterR2-selective neurons significantly increased their activity (**Fig. 3a**, solid trace; **Extended Data Fig. 6c**). Conversely, when the same D4 stimulus followed a R8 sequence (a smaller-than-expected violation), D4afterR8-selective neurons significantly decreased their activity (**Fig. 3b**, dashed trace; **Extended Data Fig. 6d**). Critically, the same D4-selective PE neurons selected from one context were modulated in the opposite direction in the inverse context. Thus, D4afterR2-selective PE neurons showed reduced activity to a smaller-than-expected D4 (after R8, **Fig. 3a**, dashed trace), and D4afterR8-selective neurons showed enhanced activity to the larger-than-expected D4 (after R2, **Fig. 3b**, solid trace). The fact that the same neurons are bidirectionally modulated despite responding to the exact same sensory input (Amplitude 4) under the same probability context, reinforces the notion that the prediction error encoded by these neurons is determined by the sign of the discrepancy relative to the expectation, not by the absolute sensory amplitude of the stimulus.

**Fig. 3.**
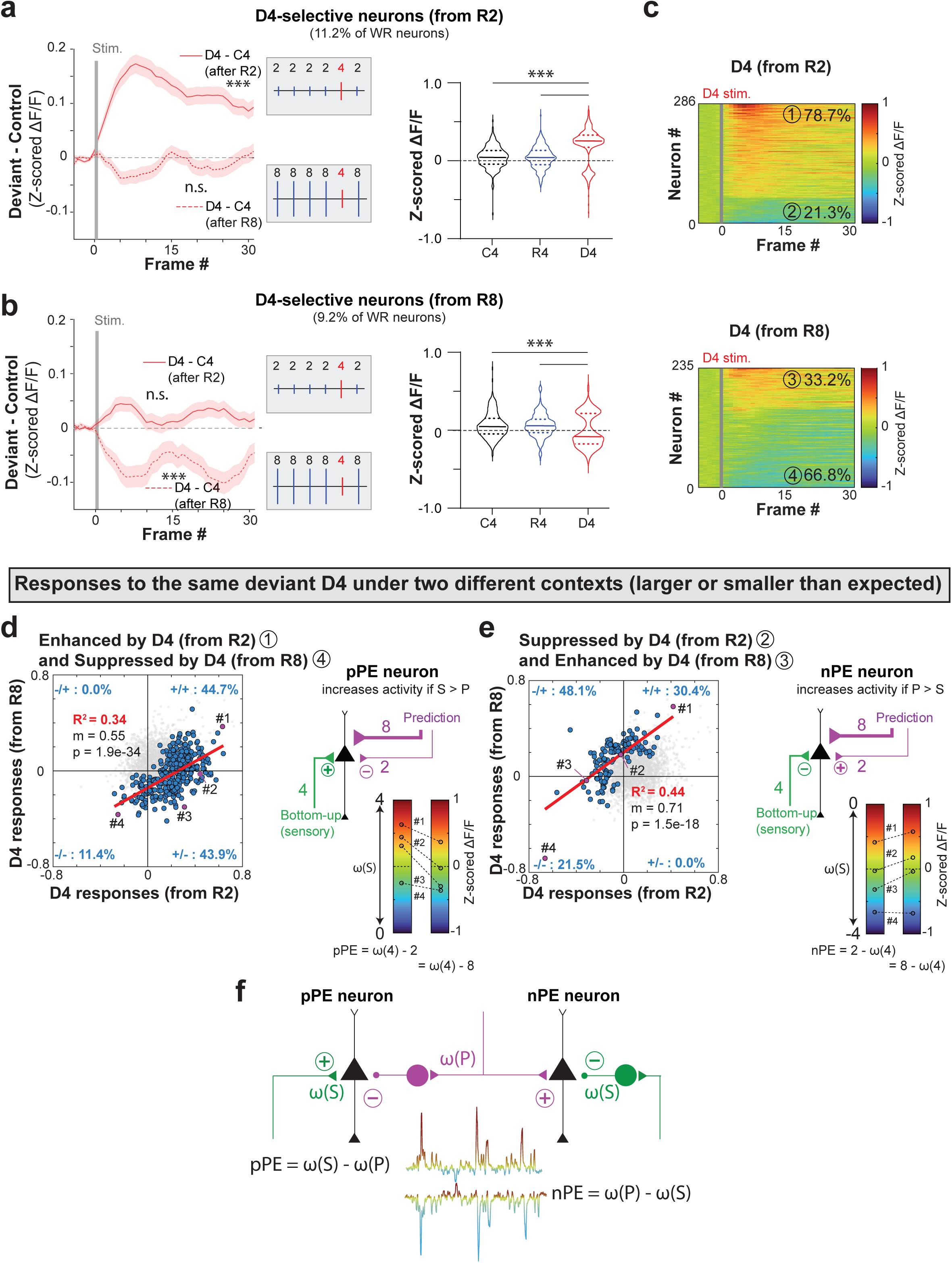
Shared prediction but distinct sensory inputs define pPE and nPE functional ensembles. **a)** Left, Subtracted mean evoked PE activity (D4 – C4, z-scored ΔF/F) of D4-selective neurons chosen in a greater-than-expected context (after R2) to a D4 stimulus (D4 – C4) in either R2 (solid line) or R8 (dashed line) contexts; Right, statistical comparison per neuron of the mean evoked activity (Linear Mixed-Effects Model, Context p<0.001, Amplitude p<0.001, Context x Amplitude interaction p<0.001, n= 286 neurons from 6 mice). **b)** Same as *a* but for D4-selective neurons chosen in a smaller-than-expected context (after R8) (Linear Mixed-Effects Model, Context p<0.001, Amplitude p<0.001, Context x Amplitude interaction p= 0.001, n= 235 neurons from 6 mice). **c)** Single-neuron mean evoked activity (z-scored ΔF/F) of D4-selective neurons after R2 (top, ① enhanced group, ② suppressed group) or after R8 (bottom, ③ enhanced group, ④ suppressed group) in response to D4, sorted by max value. **d-e)** Cross-deviant correlation scatter plots for pPE and nPE ensembles, respectively. Percentages indicate the relative distribution of neurons across quadrants. Schematic diagrams illustrating the proposed arithmetic framework for pPE and nPE neurons. Heatmap of mean evoked activity from example neurons shown in purple in panels *d* and *e*. **f)** Schematic diagram of the proposed canonical cortical microcircuit model with mirrored, asymmetrical connectivity pathways between pPE and nPE populations. ***p<0.001. Shaded areas and error bars in *a-b* represent SEM.

This conclusion was further supported by assessing the cross-context generalization of our SVM classifiers. When decoders trained to classify a D4 stimulus in one oddball paradigm were tested on the D4 responses from the other oddball, the stimulus was not identified as a D4 (**Extended Data Fig. 6e**). Notably, the cross-classification accuracy for the ‘deviant’ identity was consistently below statistical chance, underscoring the notion that, despite being the same sensory input, the internal predictive state reorganizes the neuronal representation in such a way that the same sensory stimuli in different contexts are essentially treated as distinct functional entities by the circuit.

As in the previous analysis with deviants of different amplitudes, individual neurons selectively responding to D4 were composed of two well-differentiated populations of neurons with “enhanced” or “suppressed” activity (**Fig. 3c**). Again, D4-selective neurons in each context showed very low correlations (R^2^= 0.04 and 0.003 for D4-selective neurons after R2 [groups ① and ②] and R8 [groups ③ and ④], respectively; data not shown). Grouping neurons based on their modulation polarity (either always enhanced [groups ① and ③] or always suppressed [groups ② and ④] by both D4) also showed low and negative correlations (R^2^ = 0.14 and 0.01, data not shown). But when we grouped neurons based on the majority ensemble from each group, we observed a robust positive correlation between neurons enhanced by D4 in a R2 context [group ①] and those suppressed by D4 in a R8 context [group ④], and vice versa (groups ② and ③) (R^2^= 0.34 and 0.44, respectively; **Fig. 3d-e**). This confirms a unified functional identity for both pPE neurons, which increase their activity when sensory input exceeds prediction (S > P), and nPE neurons that increase their activity when prediction exceeds sensory input (P > S).

Conveniently, our protocol also allowed us to investigate the circuit logic when the internal prediction was held constant (R4) while the direction of the sensory violation differed (D2 vs. D8). Surprisingly, even though the sensory input was different between the two contexts, grouping neurons by their functional identity as pPE or nPE (**Extended Data Fig. 6f**, left and right, respectively) neurons yielded a similar strong positive correlation (R^2^= 0.30 and 0.19, respectively). More importantly, in this case grouping by response polarity (i.e., enhanced or suppressed for both deviants) yielded a similar but negative correlation (R^2^= 0.27 and 0.16, respectively; **Extended Data Fig. 6g**), such that neurons with the greatest evoked enhancement for D8 displayed the strongest suppression for D2, and vice versa.

These correlations can be parsimoniously explained by the distribution of synaptic weights, ω (**Fig. 3d-e and Extended Data Fig. 6f-g**; example neurons), or recruitment probability, where neurons with the stronger inputs for a particular amplitude will be more strongly modulated depending on the comparison between sensory and predicted inputs. Under this framework, the relative ‘error-coding’ rank of an individual neuron remains consistent across contexts.

Under the same sensory input (D4) presented across different predictive contexts (R2 or R8, **Fig. 3d-e**), the PE for a Deviant stimulus can be expressed as:

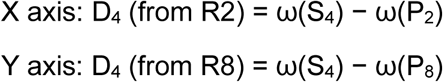

Assuming P2 and P8 are independent, the observed correlations (R^2^= 0.34 and 0.44) are driven by the covariance of ω(S_4_), i.e., the synaptic weight of individual neurons to Amplitude 4.

Conversely, when the internal prediction is held constant (R4) across varying sensory violations (D2 or D8, **Extended Data Fig. 6f-g**), the responses can be expressed as:

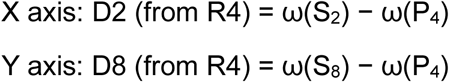

Because sensory inputs S2 and S8 drive largely non-overlapping ensembles in the Control context, the observed correlation (R^2^= 0.30, 0.19, 0.27 and 0.16) is driven instead by the covariance of ω(P_4_), i.e., the shared predictive weight for Amplitude 4.

To account for all these observations, one can represent the circuit architecture as mirrored asymmetrical connectivity pathways between pPE and nPE ensembles (**Fig. 3f**). Based on our functional correlation analysis across two complementary axes, bottom-up input engages distinct functional connectivity pathways (**Fig. 3d-e**), in contrast, the internal prediction signal is shared between both ensembles (**Extended Data Fig. 6f-g**) and can drive strong correlations independent of the different sensory inputs.

### Confirmation of the canonical microcircuit for signed prediction error

To empirically test our proposed microcircuit model, we systematically compared a fixed stimulus amplitude (4 or 8) across a wide range of contexts of higher and lower amplitudes (2, 4, 6, 8, and 10). In agreement with our model, the evoked activity of PE neurons scaled largely proportionally with the mathematical difference between the sensory input and the internal prediction (**Fig. 4a**). There was an expected sign inversion when the deviant switched from being ‘larger-than-expected’ (enhanced activity) to ‘smaller-than-expected’ (suppressed activity) (**Fig. 4b**). Furthermore, both the percentage of PE neurons amongst WR neurons and the relative proportion of pPE to nPE neurons were determined by the sign and magnitude of this difference (**Fig. 4c**). These results were observed for both fixed amplitudes (4 and 8) with remarkably similar values, in such a way that when we averaged the responses according to the signed difference (e.g., D8 - R2 and D10 - R4 = +6, etc.) a clear bidirectional modulation emerged (**Fig. 4d-e**). This demonstrates that the evoked response to an unexpected event is mainly modulated by the difference between the expected and the actual input rather than the sensory input alone.

**Fig. 4.**
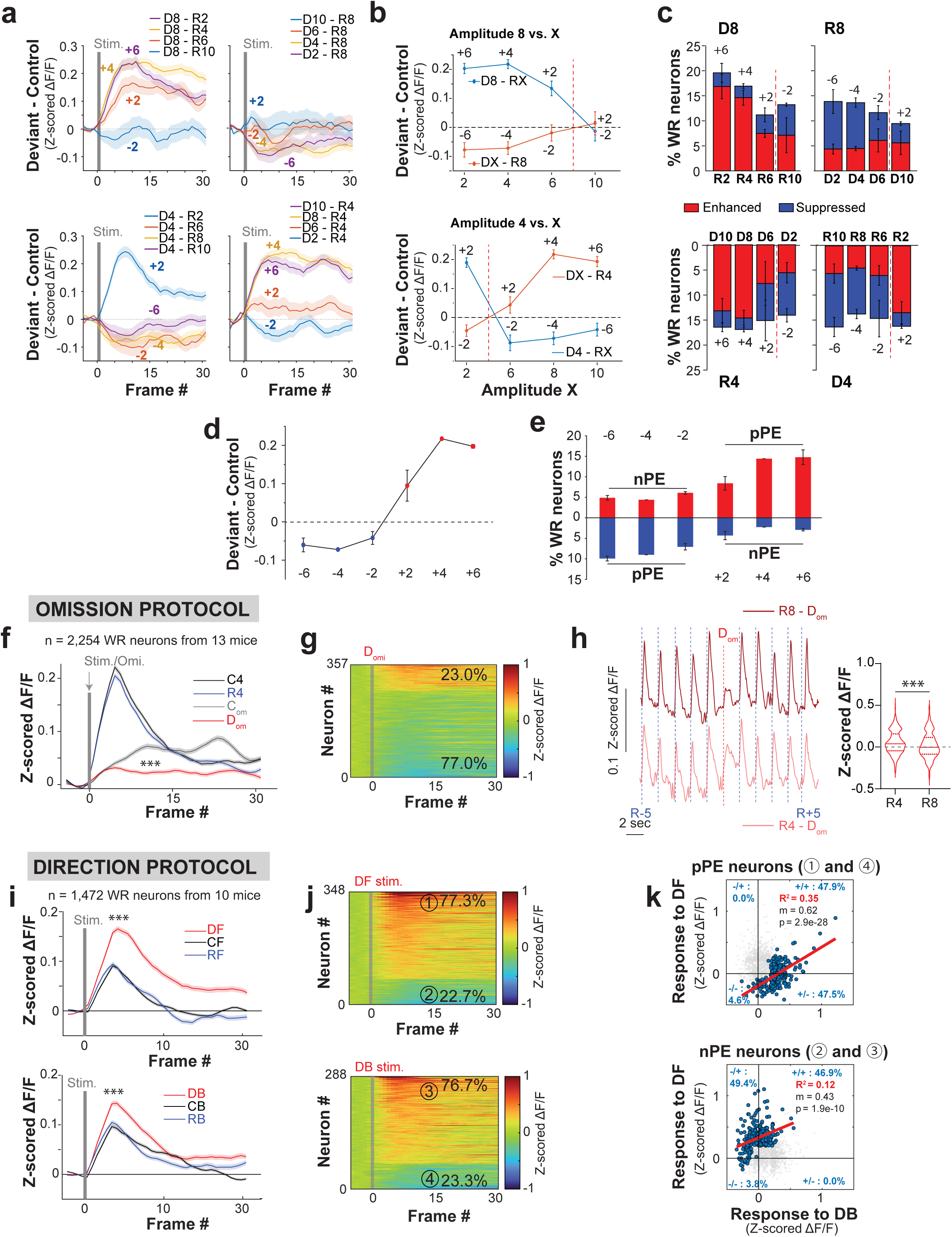
Canonical microcircuit for signed prediction in omission and direction oddball protocols. **a)** Subtracted mean evoked PE activity (D – C, z-scored ΔF/F) of Deviant-selective neurons for each amplitude comparison when D8 is fixed (top left), R8 is fixed (top right), D4 is fixed (bottom left), and R4 is fixed (bottom right). **b)** Mean evoked PE activity comparison (D – C) of Deviant-selective neurons for each amplitude comparison when Amplitude 8 (top) or 4 (bottom) is fixed. **c)** Comparison of enhanced and suppressed Deviant-selective neurons per mouse as a percentage of WR neurons for each amplitude comparison when D8 is fixed (top left), R8 is fixed (top right), R4 is fixed (bottom left), and D4 is fixed (bottom right). **d)** Averaged mean evoked PE activity (D – C) based on the signed difference. **e)** Averaged enhanced and suppressed Deviant-selective neurons per mouse as a percentage of WR neurons based on the signed difference. **f)** Mean evoked population activity of all WR neurons (n= 2,254 neurons from 13 mice) during the unexpected sensory omission protocol: Control Stimulus (C4, black), Repetitive Stimulus (R4, blue), Control Omission (C_om_, grey), and Deviant Omission (D_om_, red) (Linear Mixed-Effects Model, Context p<0.001, n= 2,254 neurons from 13 mice). **g)** Single-neuron mean evoked activity (z-scored ΔF/F) of D_om_-selective neurons in response to D_om_, sorted by max value. **h)** *Left*, Mean evoked population activity (z-scored ΔF/F) of all WR neurons during the omission of R8 (n = 489 neurons from 4 mice) and omission of R4 (n= 424 neurons from 4 mice) showing the preceding and subsequent actual stimuli; *Right*, corresponding statistical comparison per neuron (Linear Mixed-Effects Model, Context p= 0.001, N= 4 mice). **i)** Mean evoked population activity (z-scored ΔF/F) of all WR neurons (n= 1,472 neurons from 10 mice) for Forward (F, left, ① enhanced group, ② suppressed group) and Backward (B, right, ③ enhanced group, ④ suppressed group) directions across three predictability contexts (Linear Mixed-Effects Model, Context p < 0.001, Direction p= 0.468, Context x Direction interaction p<0.001, n= 1,472 neurons from 10 mice). **j)** Single-neuron mean evoked activity (z-scored ΔF/F) of DF- (top) and DB-selective (bottom) neurons in response to DF and DB, respectively, sorted by max value. **k)** Cross-deviant correlation scatter plot between different whisker deflection directions (DF vs. DB) for putative pPE and nPE neurons. Percentages indicate the relative distribution of neurons across quadrants. ***p<0.001. Shaded areas and error bars represent SEM.

We also recorded responses to a complete omission of an expected sensory input, which according to our model should generate a neuronal response representing the pure prediction signal. In this new “Oddball Omission” protocol the Deviant event was the omission (D_om_) of a predictable stimulus of amplitude 4, R4 (**Extended Data Fig. 7a**), as well as “Control Omissions” (C_om_) occurring within a randomized amplitude context. At the population level (n= 2,254 WR neurons, N= 13 mice), a non-zero number of L2/3 neurons were recruited by the D_om_ (**Extended Data Fig. 7b**), and their evoked responses were significantly smaller than for C_om_ (**Fig. 4f**; **Extended Data Fig. 7c-d**), as a result of a decrease in responsivity rather than in maximum response (**Extended Data Fig. 7e-f**). According to our model, a sensory omission (S= 0) should recruit two distinct populations of D_om_-responsive neurons: a larger subpopulation of pPE neurons that exhibit suppressed evoked activity due to the presence of a prediction without a corresponding sensory input (0 - P), and a smaller subpopulation of nPE neurons that exhibit enhancement as they signal the absence of the predicted input (P - 0). Indeed, we identified these specialized sub-populations, with proportions remarkably similar to those observed in our amplitude oddball experiments (77% pPE and 23% nPE; **Fig. 4g**).

If these neuronal responses represent the internal prediction, the omission signal should scale with the magnitude of the prediction. Accordingly, we observed that the D_om_-evoked response during a R8 context was significantly smaller (**Fig. 4h**) and a trend towards a higher proportion of D_om_-selective neurons was recruited (**Extended Data Fig. 7g**, left). Finally, by increasing the probability of the omission from 10% to 50% (random), we observed a gradual decay in the recruitment of D_om_-selective neurons (**Extended Data Fig. 7g**, right), mainly driven by a decay in the number of putative nPE neurons (enhanced neurons). To control for sensory-evoked responses driving the identification of suppressed neurons, we implemented a more stringent criteria by comparing only the evoked responses during D_om_ with C_om_, observing a dramatic decrease at 25% omission probability, and no neurons being recruited at 50% probability (**Extended Data Fig. 7h**).

Finally, to further test if this model could be extended to other stimulus features, we implemented an oddball paradigm where the Deviant was a whisker deflection in a different direction (forward/anterior vs. backward/posterior (Musall et al., 2017)) but of equal amplitude (**Extended Data Fig. 7i**). In this case, the deviant stimulus, regardless of direction, consistently evoked a larger response than in the Control context (**Fig. 4i; Extended Data Fig. 7j-l**), mainly due to an increase in responsivity (**Extended Data Fig. 7m**-**n**). We found similar proportions of enhanced (77.3% and 76.7% of Deviant-selective neurons for DF and DB, respectively) and suppressed neurons (22.7% and 23.3% of Deviant-selective neurons for DF and DB, respectively) (**Fig. 4j**), just as in the previous amplitude experiments. We also found high positive cross-deviant correlations when the enhanced and suppressed neurons from the different deviants were grouped together (**Fig. 4k**), but negative values when grouped by their modulation polarity (**Extended Data Fig. 7o**), aligning with our hypothesis of two different functional ensembles.

The observed consistency across different stimulus features suggests a common computational principle: mirrored pPE and nPE ensembles compare their sensory inputs against a shared internal prediction, generating PE signals proportional to the dissimilarity of the input synaptic weights.

### Cortical Hierarchy of Signed Prediction Errors

Having established a functional model of signed PE within L2/3 pyramidal neurons, we next sought to identify the remaining essential components of the circuit mediating these computations: the bottom-up sensory signal (relayed via the thalamus) and the top-down signal providing the prediction (the internal model based on repetition).

We first considered bottom-up inputs to vS1, which include both lemniscal, originating from the ventro-posteromedial thalamic nucleus (VPM), and paralemniscal pathways, arising from the POm (Wimmer et al., 2010). VPM and POm projections convey precise single-whisker sensory signal vs. multi-whisker contextual information about those inputs, respectively (Staiger and Petersen, 2020). We first recorded L4 spiny stellate (SS) neurons in transgenic Scnn1a-Cre;jGCaMP8s mice. L4-SS neurons are the primary source of sensory input to L2/3 pyramidal neurons that is relayed onto vS1 from VPM. In the control stimulation protocol, L4-SS neurons increased their mean activity to deflections of larger amplitudes and displayed sharp amplitude-tuning curves (**Extended Data Fig. 8a-c**), similar to L2/3 pyramidal neurons. However, during oddball protocols, instead of computing signed PE, Deviant-selective L4-SS neurons displayed only a significant increase for D8 (**Fig. 5a)**. We then recorded from axons originating in the POm after stereotaxic injections of pGP-AAV-syn-jGCaMP8s-WPRE (**Extended Data Fig. 8d**). These axons displayed little stimulus amplitude selectivity during the Control Block (**Extended Data Fig. 8e-f**) and behaved similar to L4-SS neurons during oddball protocols, showing only a significant enhancement for D8 (**Fig. 5b**). This unspecific enhancement of L4-SS neurons and POm axons suggests they operate primarily as unsigned novelty or deviant detectors.

**Fig. 5.**
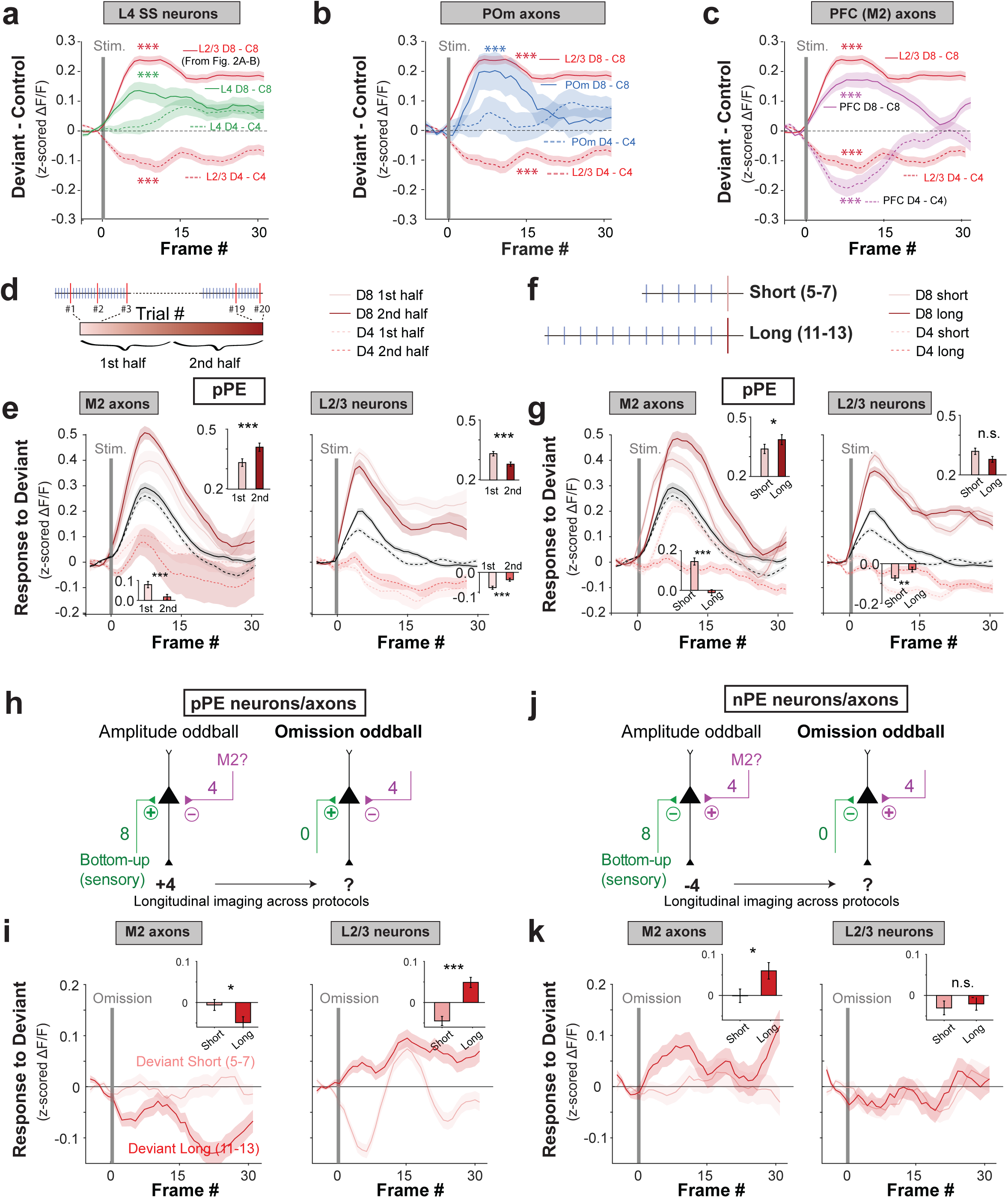
Cortical hierarchy of signed prediction errors. **a)** Subtracted mean evoked PE activity (D – C, z-scored ΔF/F) of Deviant-selective L4 spiny stellate (SS) neurons to greater- and smaller-than-expected Deviants (Linear Mixed-Effects Model, Amplitude 8: Context p<0.001, Amplitude p<0.001, Context x Amplitude interaction p<0.001, n= 148 neurons from 7 mice; Amplitude 4: Context p= 0.757, Amplitude p<0.001, Context x Amplitude interaction p= 0.075, n= 166 neurons from 7 mice). Equivalent data from L2/3 neurons from Fig. 2a-b shown for comparison. **b)** Subtracted mean evoked PE activity (D – C) for Deviant-selective POm→S1 boutons (Linear Mixed-Effects Model, Amplitude 8: Context p= 0.009, Amplitude p= 0.75, Context x Amplitude interaction p < 0.001, n = 46 boutons from 4 mice; Amplitude 4: Context p= 0.626, Amplitude p= 0.069, Context x Amplitude interaction p= 0.506, n= 58 boutons from 4 mice). **c)** Subtracted mean evoked PE activity of M2→S1 axon boutons (Linear Mixed-Effects Model, Amplitude 8: Context p<0.001, Amplitude p<0.001, Context x Amplitude interaction p<0.001, n= 202 boutons from 6 mice; Amplitude 4: Context p < 0.001, Amplitude p= 0.005, Context x Amplitude interaction p<0.001, n= 131 boutons from 6 mice). **d)** Schematic of trial-by-trial comparison between the first and second half of Deviant presentation trials. **e)** Left, Trial-by-trial mean evoked activity (z-scored ΔF/F) for the first half Deviant trials and statistical comparison (insets) for D8 (dark maroon) and D4 (light maroon) in putative pPE M2→S1 axon boutons (Amplitude 8: Paired t-test, p<0.001; Amplitude 4: Paired t-test, p<0.001, n= 232 from 6 mice); Right, same but for L2/3 pyramidal neurons (Amplitude 8: Wilcoxon test, p<0.001; Amplitude 4: Paired t-test, p<0.001, n= 445 from 10 mice). **f)** Schematic of short (5–7) and long (11–13) repetitive sequences preceding the presentation of the Deviant. **g)** Left, Short-term experience-dependent mean evoked activity (z-scored ΔF/F) and statistical comparison (insets) for D8 (dark maroon) and D4 (light maroon) in putative pPE M2→S1 boutons (Amplitude 8: Paired t-test, p= 0.031; Amplitude 4: Paired t-test, p<0.001, n = 232 from 6 mice); Right, same but for L2/3 pyramidal neurons (Amplitude 8: Wilcoxon test, p= 0.244; Amplitude 4: Paired t-test, p= 0.007, n= 445 from 10 mice). **h-k)** Schematic of longitudinal recording for putative pPE (*h*) and nPE (*j*) ensembles across amplitude and omission oddball blocks. Mean evoked activity and statistical comparison (insets) for Long (dark red) and Short (light red) repetitive sequences preceding a deviant omission in putative pPE (*i*) M2→S1 axons (left; Paired t-test, p= 0.034, n= 282 from 5 mice) and L2/3 pyramidal neurons (right; Paired t-test, p<0.001, n= 413 from 6 mice), and nPE (*k*) M2-to-S1 axons (left; Paired t-test, p= 0.026, n= 141 from 5 mice) and L2/3 pyramidal neurons (right; Paired t-test, p= 0.683, n= 212 from 6 mice). *p<0.05; **p<0.01; ***p<0.001. Shaded areas and error bars represent SEM.

Lasty, we examined the functional properties of top-down projections by recording from prefrontal axons after stereotaxic injections of pGP-AAV-syn-jGCaMP8s-WPRE in the secondary motor cortex (M2; **Extended Data Fig. 8h**). These axons displayed an intermediate level of feature selectivity to control stimuli (**Extended Data Fig. 8i-j**). In the oddball protocol, unlike L4-SS neurons and POm axons, Deviant-selective M2→S1 boutons precisely recapitulated the bidirectional, signed modulation observed in L2/3 neurons (**Fig. 5c; Extended Data Fig. 8k**). The predictive processing framework posits that these top-down projections must carry additional information related to the certainty of the prediction (Friston, 2018; Hodson et al., 2024). To test this, we compared the responses of M2→S1 axons and L2/3 neurons with respect to how evidence accumulation modulated their signed PE activity. First, we analyzed the trial-by-trial evolution of their PE activity, by comparing the average value of the first half of the Deviant trials, when they are most novel, to the second half of the trials, when the animal has already experienced many unexpected stimuli (**Fig. 5d**). We found that M2→S1 axons undergo a bidirectional experience-dependent amplification of their error signals, such that, as more deviants are encountered, pPE axons magnify their signed PE divergence with a progressive increase in responses to the larger-than-expected D8 and a decrease in responses to the smaller-than-expected D4 (**Fig. 5e**, left). In contrast, pPE L2/3 neurons show the opposite modulation, a progressive attenuation of their bidirectional PE activity (**Fig. 5e**, right). Thus, as the Deviant becomes more familiar, the prediction signal from prefrontal regions gets stronger, while in vS1, the strength of prediction errors lessens, at least for pPE neurons (the difference between nPE M2→S1 axons and nPE L2/3 neurons was less striking; **Extended Data Fig. 8l**).

Second, we evaluated the differences in PE activity according to a shorter-term internal model confidence, by comparing neural responses to a Deviant depending on how many previous repetitive stimuli preceded it. Although our experimental protocol was not specifically optimized for this analysis (the only condition in our protocol was a minimum of 5 repetitions before the deviant; see Methods), we split Deviant presentations into those that followed short or long repetition blocks (5–7 vs.11-13, respectively; **Fig. 5f**). For M2→S1 pPE axons, as the number of repetitions increased, the difference in signed PE activity was again magnified (**Fig. 5g**, left). In contrast, L2/3 pPE neurons showed the opposite modulation: the higher the number of previous repetitions the lower the signed PE (**Fig. 5g**, right). However, we did not find any significant modulation for the nPE axons or neurons (**Extended Data Fig. 8m**), either because we were underpowered to detect any, or because perhaps nPE neurons are less influenced by experience in the shorter term. Taken together, these findings suggest that M2→S1 axons implement a highly elaborated version of signed PE, where the gain of the error signal is scaled by the confidence in the internal model, or precision (γ) (γ x |Sensory - Prediction|) of the established regularity. L2/3 pyramidal neurons, on the other hand, scale the PE signal negatively as the certainty increases (−γ x |Sensory - Prediction|).

To further test our hypothesis that pPE and nPE elements within M2→S1 axons and L2/3 neurons display different precision-dependent dynamics, we performed longitudinal imaging to track the same axons/neurons across amplitude and omission oddballs (**Fig. 5h-k**). In agreement with our signed comparator model, when we identified Deviant-selective M2→S1 axons based on their responses during the amplitude oddball we found that the pPE subset (those whose activity increased when S>P and decreased when S<P, **Fig. 5h-i**) exhibited a stronger suppression to the D_om_ after Long repetition sequences, i.e., when the internal expectation was highest, than after Short sequences (**Fig. 5i**, left). In contrast, L2/3 pPE neurons showed an increase in activity after Long repetitions (**Fig. 5i**, right). Conversely, M2→S1 nPE axons (whose activity decreases when S>P and increases when S<P, **Fig. 5j-k**) demonstrated the reciprocal functional trajectory, that is, an expected enhancement in activity during high-certainty omissions (Long sequences, **Fig. 5k**, left), while nPE L2/3 neurons showed no differences between Short and Long repetitions (**Fig. 5k**, right).

In summary, these comparative analyses reveal a distinct hierarchical processing between L2/3 pyramidal neurons and top-down M2→S1 inputs. Prefrontal feedback projections fully met all canonical criteria of predictive processing theories, exhibiting robust, bidirectional signed PE signals that are positively weighted by prediction confidence. Conversely, the PE activity of L2/3 neurons was negatively weighted by prediction confidence.

## Discussion

In this study, we demonstrate that L2/3 pyramidal neurons of vS1 simultaneously encode tactile stimulus amplitude and its statistical predictability through specialized pPE and nPE ensembles. By comparing bottom-up sensory inputs with top-down expectations, this circuit signals both larger-and smaller-than-expected whisker deflections. This computation is performed at multiple steps across the cortical hierarchy: 1. Bottom-up POm→S1 projections and L4-SS neurons operate as unsigned novelty detectors; 2. L2/3 pyramidal neurons compute a local signed PE that decreases with model certainty; and 3. Top-down PFC/M2→S1 feedback projections calculate a signed PE that scales positively with model certainty.

Could our findings be explained by a traditional SSA framework, whereby bottom-up gain modulation resets baseline neuronal firing based on repetition suppression (Adibi and Lampl, 2021)? There are several reasons why we believe classical adaptation alone cannot explain the signed responses we observed in this particular study. First, we employed a long inter-stimulus interval (2 s) relative to the much shorter whisker deflection (50 ms). This minimized the contribution of short-term synaptic depression compared to the nearly continuous stimulation protocols used in other studies where SSA is certainly likely to play a role (Adibi et al., 2013; Musall et al., 2014, 2017). Indeed, the responses between the Control and the Repetitive conditions to all amplitudes tested were very similar, indicating very little adaptation (e.g., **Figs. 1f-h**, **3a-b, 4f, 4i**). Second, passive adaptation inherently requires a physical sensory stimulus to suppress or drive neuronal activity and, therefore, SSA cannot explain the recruitment of negative prediction error (nPE) ensembles that increased their neuronal response in the total absence of bottom-up input (**Fig. 4g**). Third, SSA cannot account for the coexistence of intermingled pPE and nPE ensembles that undergo simultaneous opposite modulations to the exact same physical stimulus (**Fig. 3c**). Finally, SSA cannot explain how M2 feedback projections progressively amplify signed error signals as the expectation consolidates, a gain-scaling trajectory that is exactly the opposite of adaptation.

In fact, the simultaneous encoding in L2/3 neurons of stimulus amplitude and predictability that we report can reconcile classical SSA frameworks and the feature-selective sparse coding observed in vS1 with dynamic predictive modulation. On one hand, during the predictively neutral control condition, a small percentage of L2/3 pyramidal neurons were whisker-responsive and highly selective for specific stimulus amplitudes. This sparse and robust ensemble-based coding (Petersen et al., 2003; Gollnick et al., 2016) is consistent with the classical ‘iceberg effect’ (Adesnik and Scanziani, 2010; Voelcker et al., 2022), wherein a sparse subset of pyramidal neurons emerges above a local inhibitory threshold. Under this bottom-up scheme, feedforward sensory inputs simultaneously excite feature-tuned pyramidal neurons and broadly tuned GABAergic interneurons, establishing a generalized inhibitory background that permits only the most strongly and specifically driven pyramidal neurons to reach the action potential threshold (Kerlin et al., 2010; Atallah et al., 2012; Petersen and Crochet, 2013). On the other hand, our findings reveal a second mechanism that dynamically adjusts the height of the iceberg. The prediction of the incoming stimulus amplitude establishes a subtractive template against which incoming sensory inputs are compared, such that the absolute difference between these two inputs determines the height of the iceberg, while the algebraic sign of this difference dictates the proportion of the ensemble positioned above or below it (**Extended Data Fig. 9**). Consequently, when the stimulus amplitude exceeds internal expectations (S > P; e.g., 8 - 4 = +4), evoked activity increases, recruiting a larger proportion of pyramidal neurons above the inhibitory threshold. Conversely, when the stimulus amplitude falls below internal expectations (S < P; e.g., 4 - 8 = −4), the subtractive template drives suppression, effectively decreasing the population response below the ‘waterline’ set by the expected amplitude. This circuit architecture allows L2/3 to maintain its high feature specificity while ensuring its output is continuously normalized (Carandini and Heeger, 2012) and contextualized by an internal model of the sensory environment.

Within this framework, we identified two mirroring ensembles that closely align with previously proposed pPE and nPE ensembles (Keller and Mrsic-Flogel, 2018). Because pyramidal neurons produce non-negative outputs at exceptionally low baseline firing rates, a single error-encoding population could not signal deviations in both directions and, hence, dedicated nPE neurons must exist to selectively signal when a stimulus falls short of expectations. While recent studies have begun to uncover evidence of nPE-like activity, these empirical observations have largely relied on sensory omissions or complex, multimodal sensorimotor mismatches (Audette and Schneider, 2023; Lao-Rodríguez et al., 2023; O’Toole et al., 2023; Leonardon et al., 2025; Solyga and Keller, 2025; Yaron et al., 2025), limiting our ability to precisely manipulate and isolate the internal expectation. Our oddball paradigm has the advantage that it isolates sensory inputs from internally generated expectations independently, which allowed us to identify both error ensembles within the same local circuit. Crucially, both pPE and nPE ensembles displayed increases but also decreases in their evoked activity in comparison to the same stimulus in a predictively neutral context. Furthermore, this modulation was not restricted to stimulus amplitude but seems to be based on the synaptic input dissimilarity. Across a wide range of stimulus features (five amplitudes, opposite directions, and omissions), the greater the distance between the bottom-up input and the prediction, the larger the PE response, and the greater the proportion of recruited pPE relative to nPE neurons (quite similar values to previous studies (O’Toole et al., 2023)). The simplicity of this mechanism underscores its utility as a potential canonical computation beyond vS1 and irrespective of the nature of the sensory input, as future studies may reveal.

As originally proposed (Keller and Mrsic-Flogel, 2018), the canonical prediction circuit (**Fig. 3f**) must include inhibitory neurons that help implement the signed response of pPE and nPE neurons. Previous studies demonstrated that L4 amplifies the feedforward sensory representation of unexpected stimuli transmitted to superficial layers (Yu et al., 2016) through connections not just with pyramidal neurons but also with parvalbumin (PV) interneurons, in reciprocally interconnected local subnetworks (Goz and Hooks, 2023). At the same time, POm is recruited during sensory novelty (Audette et al., 2019; Zhang and Bruno, 2019) and directly activates PV interneurons but also vasoactive intestinal polypeptide interneurons (Audette et al., 2018; Sermet et al., 2019; Williams and Holtmaat, 2019), pointing to a potential gating mechanism. Clarifying the precise role of distinct interneuron classes in deviance detection will be necessary to complete the microcircuit (Jamali et al., 2024; Ding et al., 2026).

The functional nature of the PFC/M2→S1 projection represents a key concept in hierarchical predictive processing. Although some evidence has pointed to the relevance of prefrontal inputs for the generation of PE in primary sensory areas (Banerjee et al., 2020; Hamm et al., 2021; Bastos et al., 2023; Hockley et al., 2025; Tsukano et al., 2026), the precise computational variable they transmit remains disputed (Gabhart et al., 2025). We show that M2→S1 feedback axons recapitulate signed PE dynamics, consistent with recent theories stating that genuine error computations emerge prominently in frontal networks and are sent back to primary cortices (Banerjee et al., 2020; Gabhart et al., 2025). Remarkably, our longitudinal tracking of putative pPE and nPE axons during unexpected sensory omissions suggests that PFC/M2 feedback – most probably L5 intratelencephalic (IT) neurons – supplies the internal prediction template back to S1 under high certainty. This aligns with theoretical frameworks proposing that top-down expectations are formed by the linear accumulation of PEs over time (Bastos et al., 2012). Under this predictive model, the progressive increase of signed errors observed in M2→S1 axons during rising certainty may reflect the active integration of vS1-PE signals, allowing L5-IT neurons in PFC/M2 to continuously build and update internal representations.

Together, these results provide direct physiological evidence of a bidirectional predictive mechanism in the cerebral cortex, in line with similar predictive systems within the cerebellum (Wolpert et al., 1998; Ohmae and Medina, 2015; Kakei et al., 2026) and the basal ganglia (Schultz et al., 1997; Bogacz, 2020; Bastos et al., 2023; Dudhabhate and Costa, 2026).

## Acknowledgements

We thank Dean Buonomano, Dario Ringach, Saray Soldado-Magraner, William Zeiger and Matteo Mariani for helpful discussions about this project. We used generative AI models to write some of the early drafts of the manuscript (Main text and figure legends), but this was subsequently thoroughly edited by the authors. We also used Gemini (3.1 pro model, Google) to generate code to analyze some of the calcium imaging data. Code was then independently inspected, edited and validated by C.A.S.L. This work was supported by the National Institute of Neurological Disorders and Stroke (R01NS117597 to C.P.-C.), National Institute of Child Health and Development (R01HD108370 and R01HD054453 to C.P.-C).

## METHODS

### Mouse lines

All experiments followed the U.S. National Institutes of Health guidelines for animal research, under an animal use protocol (ARC#2007-035) approved by the Chancellor’s Animal Research Committee and Office for Animal Research Oversight at the University of California, Los Angeles. Mice were housed in a vivarium with a 12/12 h light/dark cycle and experiments were performed during the dark cycle. Adult mice (2-4 months) of both sexes were used in this study.

To obtain Cre-dependent expression of jGCaMP8s in pyramidal neurons we crossed the reporter line TIGRE2-jGCaMP8s-IRES-tTA2-WPRE (JAX strain # 037952) with Slc17a7-Cre mice (vesicular glutamate transporter 1-expressing cells; JAX strain # 023527). To image L4 spiny stellate neurons we crossed Scnn1a-Cre mice (JAX strain # 009613) to the Ai162 (GCaMP6s) reporter line (JAX strain # 031562). For calcium imaging of axon boutons from POm or M2 (see below for coordinates), we injected WT mice (JAX strain # 000664) with an AAV1 (pGP-AAV-syn-jGCaMP8s-WPRE; Addgene # 162374-AAV1).

### Cranial window surgery and viral injections

Cranial window surgery was performed on mice at P45-P90, as described previously (Mostany and Portera-Cailliau, 2008; Holtmaat et al., 2009). Mice were anesthetized with isoflurane (5% induction, 1.5-2% maintenance via a nose cone) and placed in a stereotaxic frame (Kopf). Carprofen (5 mg/kg, i.p., Zoetis) and dexamethasone (0.2 mg/kg, i.p., Vet One) were provided for pain relief and mitigation of edema, respectively. Under sterile conditions, a 4 mm diameter craniotomy was performed centered above the right vS1 (AP = - 0.6 mm; ML = + 3 mm; relative to Bregma (Paxinos and Franklin, 2019) taking care to not damage the dura. The craniotomy was then covered with a 4 mm glass coverslip and secured by cyanoacrylate glue and dental cement. Animals with extensive subdural hemorrhages or in which the dura was punctured were discarded (Grienberger et al., 2022).

For prefrontal virus injections, we performed a 1 mm craniotomy over M2 (AP = + 1 mm; L = -+ 0.5 mm; Depth = −0.3 to −1.5 mm relative to Bregma) at the time of the cranial window surgery. Virus injections into POm were done through the open 4 mm cranial window (AP = - 2 mm; L = -+ 1.25 mm; Depth = −3.1 mm relative to bregma). Using a Picospritzer (General Valve, 30 pulses of 6 ms, 30 psi) we injected AAV1-Syn-GCaMP8s-WPRE-SV40 (ADDGENE 162374) using glass micropipettes (Sutter Instrument, 1.5 mm outer diameter, 0.86 mm inner diameter) to inject 250-350 nL. The needle was left in place for 10 min to allow for diffusion and was slowly removed to prevent backflow into the needle path. A custom horseshoe-shaped titanium head bar (3.15 mm wide × 10 mm long) was affixed to the skull with dental cement to secure the animal to the microscope stage. Animals recovered from the procedure and were fully ambulatory within 1 h after the surgery. Carprofen (5 mg/kg, s.c., once daily) was used for analgesia for 3 d post-op. We allowed 2-3 weeks for viral expression before beginning calcium imaging.

### Histology for virus injections

To confirm the correct targeting of virus injections, mice were perfused with 4% paraformaldehyde (PF) after the last recording session. Brains were harvested and post-fixed overnight in 4% PF, then stored in phosphate buffered saline with 0.02% sodium azide at 4°C. Brains were sectioned at 100 μm on a vibratome (Leica VT1000). Injection sites were then verified by taking high-resolution fluorescence microscopy images (Zeiss ApoTome2, Zen2 software; 5X objective, 0.3 NA) and aligning them to the Allen Brain Atlas.

### Intrinsic signal imaging

We mapped the location of the C2 barrel within S1 using intrinsic signal imaging, 5-7 days after cranial window surgery, as described previously (Johnston et al., 2013; Dobler et al., 2024). The cortical surface was illuminated by green LEDs (535 nm) to visualize the superficial vasculature. The macroscope was then focused ∼300 μm below the cortical surface and red LEDs (630 nm) were used to record intrinsic signals, with frames collected at 30 fps from 0.9 s before until 1.5 s after stimulation, using an 8-megapixel CCD camera (catalog #8051M-USB, Thorlabs), and custom routines written in MATLAB. Thirty trials separated by 20 s were conducted for each imaging session. The contralateral C1 whisker was gently attached with bone wax to a glass microelectrode coupled to a ceramic piezo-actuator (PI127, Physik Instrumente). Each stimulation trial consisted of a 100 Hz sawtooth stimulation lasting 1.5 s. The response signal was divided by the average baseline signal, summed for all trials, with a threshold at a fraction (65%) of maximum response to delineate the cortical representation of the stimulated whisker and guide our posterior imaging recordings.

### *In vivo* calcium imaging in head-fixed mice

In vivo 2-photon calcium imaging (Golshani and Portera-Cailliau, 2008; Mostany et al., 2015; Grienberger et al., 2022) was performed in awake mice. We used a commercial 2-photon (2P) microscope (DIY Bergamo, ThorLabs) equipped with galvo-resonant scanning mirrors, amplified non-cooled GaAsP photomultiplier tubes (Hamamatsu), a 16X objective (0.8NA, Olympus, Thor N16XLWD-PF), and ThorImage software. The microscope was coupled to a Chameleon Ultra II Ti:sapphire laser (Coherent) tuned to 930 nm, and the average power at the sample was kept <80 mW. For neuronal recordings, we recorded fluorescence activity within a single field of view (FOV) per animal (512 x 512 pixels, 834 x 834 µm) in L2/3 (200-250 µm depth) or L4 (300-400 µm depth). For axonal imaging (< 100 µm depth), we used a X2 magnification (512 x 512 pixels, 413 x 413 µm). We centered our recordings over vS1 by the location of the C2 map from the intrinsic signal imaging. Calcium imaging was performed at a framerate of 15.1 Hz. Mice were habituated to the microscope setup prior to imaging. This process lasted ∼5 days and involved a gradual progression from basic daily handling (5 min/day) until mice were comfortable with head fixation and body restraint in a plexiglass tube for periods up to 30 min. For the last 2 days of habituation, 100 trials of whisker stimulation were applied at random amplitudes.

Whisker stimulation was applied using a “comb” of von Frey Nylon filaments intercalated between whiskers (filaments were spaced 0.5 mm apart). This comb was coupled to a piezoactuator (PI127, Physik Instrumente) controlled by MATLAB to deliver the different patterns of stimulation. We recorded neuronal responses to the same physical stimulus across distinct predictability contexts in one single recording (∼20 min): 1) an ‘Oddball 1’ block in which was the Repetitive (R) or standard stimulus at 90% probability and a different stimulus was the Deviant (D) at 10% probability; 2) a flipped ‘Oddball 2’ block (R was the previous D stimulus and vice-versa). In addition, during the same imaging session, we also presented a Control (C) block in which different stimuli appeared with equal probability. This stimulation protocol allowed us to compare responses to the exact same physical stimulus (e.g., Amplitude 4) under three levels of expectation: highly predictable (Repetitive), unexpected violation (Deviant), and predictively neutral (Control). We performed three types of mismatch experiments:

Amplitude oddballs: 50 ms-long deflections of the whiskers at different amplitudes in the antero-posterior plane, followed by 1.95 s-long inter-stimulus interval.

Omission oddballs: 50 ms-long deflections of the whiskers at different amplitudes, including stimulus omissions (0 amplitude), in the antero-posterior plane, followed by 1.95 s-long inter-stimulus interval.

Direction oddballs: 50 ms-long anterior (or posterior) deflections of the whiskers at the same amplitude, followed by 1.95 s-long inter-stimulus interval during which the stimulator retracted to its initial position.

To confirm that most whiskers on one side of the snout were being deflected consistently across different bouts, we used two cameras (FLIR BFS-U3-23S3M-C: Monocamera) at 30 and 90 fps to record both the position of the comb of Nylon filaments relative to the whiskers and body movements (pupil diameter and mouse whiskers). Displacements of the comb were measured using DeepLabCut (resnet 50, shuffle1, iterations 30000) (Extended Data Fig. 1b). To train the network, two markers were placed at the end of two filaments, and two markers with a separation of 1 mm between them in a scale placed behind the filaments. After training, the positioning of the markers was exported and analyzed on Python 3. Markers with a likelihood value less than 90 were excluded and the nearest point with a likelihood greater than 90 was substituted instead. Distance of deflections per marker was calculated for each video frame and values were Z-score normalized.

### Initial processing of calcium imaging data

Calcium imaging data analysis was done with custom-written MATLAB routines (MATLAB version 2020a). Motion correction and ROI segmentation were done using Suite2p (Pachitariu et al., 2017). For neuronal detection, automated ROI exclusion was performed using a custom classifier built using previous data, followed by a manual refinement step to include or exclude ROIs that had been missed by the classifier. For analysis of axonal boutons, we used Suite2P for automated detection, followed by manual curation to confirm the ROIs bouton-by-bouton. Neuropil subtraction was performed by removing the local fluorescence signal surrounding each ROI. Fluorescent activity was normalized for each individual neuron and within each experimental block by calculating the Δ*F/F*0 signals. We estimated baseline fluorescence (*F*0) by a Gaussian mixture model with two components fitted on the raw fluorescence data. The minimum mean of the fitted Gaussian components was used as *F*0. To control for baseline drifts and equalize signal variance across distinct experimental blocks, the normalized traces were Z-scored by subtracting the mean Δ*F/F* value per block and dividing it by the standard deviation of the mean. Finally, the *z*-scored Δ*F/F* traces were temporally smoothed using a moving-average filter (5 frames) to eliminate high-frequency noise.

### Analysis of whisker responses

We averaged each neuron *z*-scored Δ*F/F* across all trials belonging to a particular stimulus (e.g. Amplitude 4) and block (e.g. Control, Deviant, Repetitive). The response to each whisker deflection was calculated using the mean *z*-scored Δ*F/F* signal averaged over a window from frames 3 to 15 (∼1 s) after stimulus onset, baseline-subtracted using the mean *z*-scored Δ*F/F* signal during the 5 frames before stimulus onset for each stimulus and block (Extended Data Fig. 1E). Neurons were classified as whisker-responsive (WR) for a particular whisker deflection amplitude if their mean response to that amplitude was larger than 0.2 *z*-scored Δ*F/F*.

We calculated a selectivity index (SI) for each neuron, defined as the response (mean *z*-scored Δ*F/F* signal averaged over a window from frames 3 to 15 after stimulus onset) to the preferred amplitude minus the sum of responses across all other amplitudes:

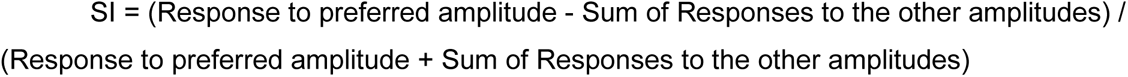

To identify individual Deviant-selective neurons that exhibited statistically significant modulation in response to sensory violations, we implemented a non-parametric Monte Carlo permutation and resampling test to generate a neuron-specific null distribution. For each recorded neuron, trial-type labels (e.g., D4, R4, C4 and D8, R8, C8) were randomly shuffled 10,000 times to construct a cell-specific null distribution representing the chance-level variance of contextual modulation. The mean response of these shuffled trials was calculated to build an empirical, neuron-specific surrogate distribution. Statistical significance thresholds were derived directly from the empirical cumulative distribution function of the surrogate data. Neurons were classified as Deviant-enhanced or Deviant-suppressed if their true Deviant response exceeded the 95 % percentile or fell below the 5 % percentile of the surrogate distribution, respectively. In the case of Omission Oddballs, an additional permutation test was performed to select neurons that exhibited statistically significant modulation only between the Devian Omission and the Control Omission.

### Statistical analyses

#### Software and statistical analysis

We used MATLAB (2023a) and ImageJ/FIJI for data processing and visualization. We used MATLAB, IBM SPSS version 25 (IBM, Armonk, NY) and GraphPad Prism 11 for statistical analyses.

For comparisons between mice, normality was assessed using the D’Agostino & Pearson test (α = 0.05). Mauchly’s test of sphericity was conducted, and the Greenhouse-Geisser correction was applied when necessary. Statistical significance was inferred using either a One-Way Repeated Measures ANOVA followed by Tukey’s post-hoc test for normally distributed data or a Friedman Repeated Measures Analysis of Variance on Ranks, followed by Dunnett’s post hoc test when the normality assumption was violated.

For comparisons between neurons, to account for the hierarchical nesting and non-independence of our imaging data (which isolates the true biological variance of individual cells from confounding animal-specific batch effects), data were analyzed using Linear Mixed-Effects Models (LMMs; MIXED procedure in SPSS). Separate and independent LMMs were constructed for three distinct dependent variables: mean calcium activity (MeanActivity), peak amplitude activity (MaxActivity), and responsivity index (ResponsivityActivity). For all models, Stimulus (e.g., Amplitude 4 vs. Amplitude 8) and Block (e.g., Control, Deviant, Repetitive), along with their crossed interaction (Stimulus x Block), were entered as fixed factor effects. Type III sum of squares was utilized to accommodate any unbalanced cell sizes across conditions. To model the multi-level dependency of the dataset, a nested random-effects covariance architecture was implemented. Specifically, a random intercept was estimated for each individual animal (Subject: MouseID) to control for inter-animal variability, and an additional random intercept was nested for individual neurons within each mouse (Subject: MouseID x NeuronID). Variance Components (VC) was selected as the covariance structure for the random effects, and parameter estimation was performed using the Restricted Maximum Likelihood (REML) method. In the event of statistically significant interactions or main effects, Estimated Marginal Means (EMMEANS) post-hoc test was applied, utilizing the Bonferroni adjustment method.

To assess history-dependent changes during the trial-by-trial evolution, we compared neuronal activity between the first and second halves of the deviant trials. Concurrently, to evaluate the impact of prior regularities, trials were categorized into two distinct, non-overlapping blocks representing low and high evidence accumulation states: Short (5–7 preceding repetitions) and Long (11–13 preceding repetitions).

Normality was assessed using the Lilliefors test (α = 0.05). Pairwise statistical significance was inferred using either a two-tailed paired Student’s t-test for normally distributed data or a Wilcoxon signed-rank test when the normality assumption was violated.

Statistical significance was set for all statistics at p < 0.05 (\**p* value < 0.05, \*\**p* value < 0.01, \*\*\**p* value < 0.001). For all analyses, the results are shown as mean ± standard error of the mean (SEM).

#### Neural decoding analyses

We decoded the specific stimulation context (e.g., Deviant, Control, Repetitive) from the neural population activity at the different aligned time points. We fitted Support Vector Machine (SVM) models (in MATLAB) with a linear kernel using the averaged neural activity within a sliding temporal window (5 frames) to classify the distinct sensory predictability conditions. For multi-class problems, the decoders distinguished between the different classes using an Error-Correcting Output Codes (ECOC) approach. We balanced the number of trials across classes by subsampling to the minimum number of trials available across conditions for each mouse.

The activity of individual neurons was standardized prior to fitting by centering and scaling the data internally within the SVM templates, which avoided data leakage from the training to the test folds. We then partitioned and evaluated the data using a 10-fold cross-validation procedure. To estimate the generalization of the decoders across different experimental paradigms, models were trained on the full training datasets and evaluated on separate test datasets using a bootstrapping approach. Specifically, test trials were randomly sampled with replacement, and this procedure was repeated across multiple iterations (200 times). The final model accuracies represent the averages over all iterations and their corresponding standard errors. As a control, in parallel we fitted models where we randomly shuffled the condition labels to establish an empirically derived chance baseline. This permutation process was repeated 1,000 times using a 5-fold cross-validation to extract the 5^th^ and 95^th^ percentiles of the shuffled null distribution. In addition to the sliding window approach, we also trained static decoders following the same procedures but averaging the neural activity across a fixed temporal window of interest (frames 3 to 20) following stimulus onset.

To analyze the similarity of the neural representations between the different classes we constructed confusion matrices. To do this, we calculated the percentage of times the cross-validated decoders correctly and incorrectly classified trials as each of the different stimulation contexts for every given true context Values are reported as the percentage of predicted trials relative to the total number of true trials for each class.

---

**Extended Data Fig. 1.**
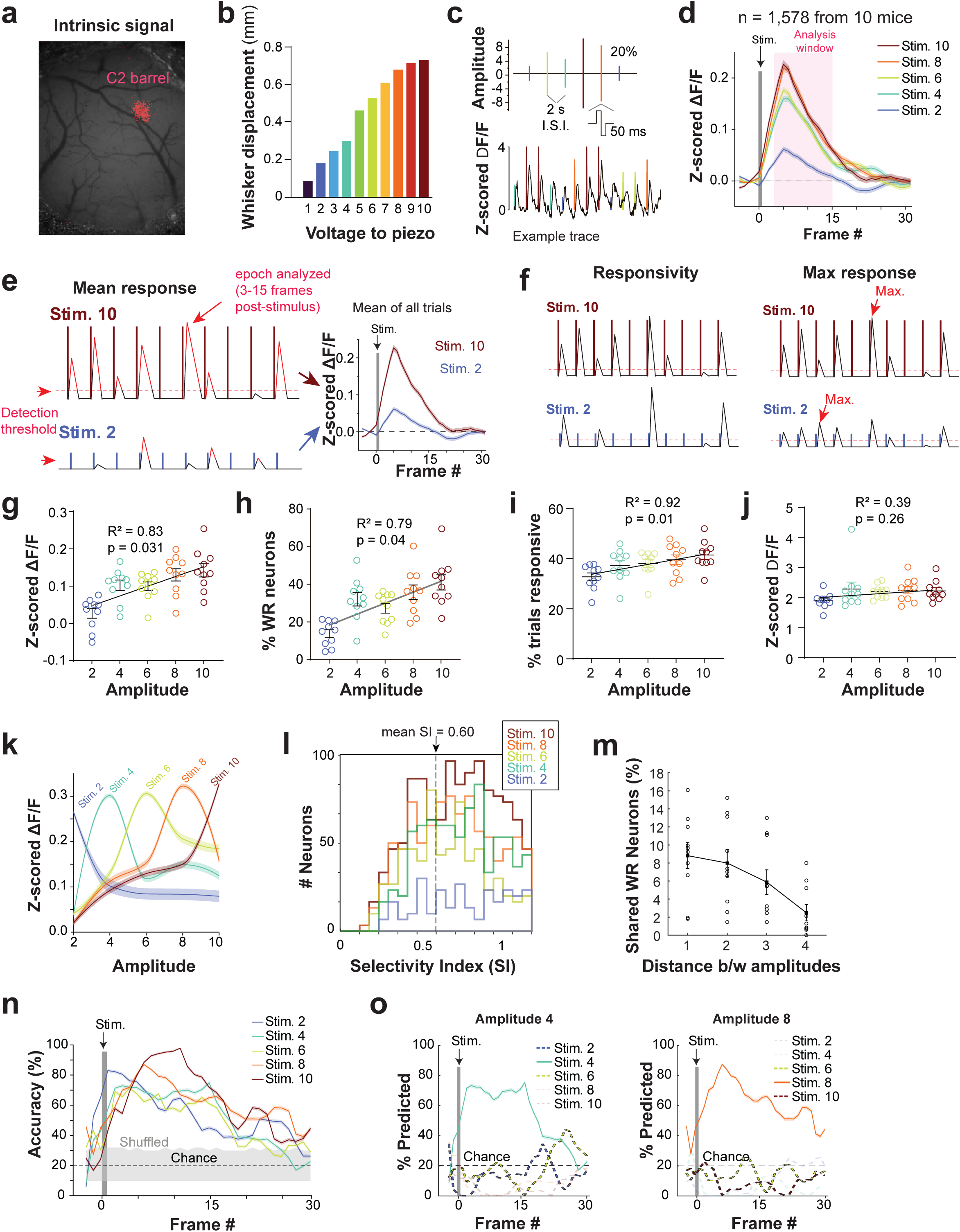
Characterization of tactile amplitude encoding in L2/3. **a)** Intrinsic signal imaging indicates C2 whisker map in vS1. **b)** Relationship between voltage sent to the piezoelectric actuator and displacement of the whisker-stimulation comb. **c)** Top, control stimulation protocol. Randomized whisker deflection amplitudes (2, 6, 8, and 10 V) delivered at regular intervals (A-P direction; square pulse, 50 ms duration, inter-stimulus interval, I.S.I. = 2 s;). Bottom, representative z-scored ΔF/F calcium trace from a representative whisker-responsive (WR) neuron. **d)** Mean evoked population activity (z-scored ΔF/F) of all WR neurons relative to all stimulation amplitudes (n= 1,578 neurons from 10 mice). The shaded pink box indicates the post-stimulus analysis window (frames 3 to 15). **e)** Schematic of mean evoked activity calculation for two example amplitudes. **f)** Left, Schematic of responsivity changes or, right, maximum evoked response driving the differences in mean evoked activity. **g)** Linear correlation between stimulus amplitude and mean evoked population activity per mouse (Simple linear regression, R^2^ = 0.833, p = 0.031, N = 10). Individual mice data represented in circles. **h)** Linear correlation between stimulus amplitude and the percentage of recruited WR neurons per mouse (Simple linear regression, R^2^ = 0.789, p = 0.044, N = 10). **i)** Linear correlation between stimulus amplitude and responsivity changes. **j)** Linear correlation between stimulus amplitude and maximum evoked response. **k)** Tuning curves of the WR neurons for different amplitudes. **l)** Distribution of Selectivity Index values (SI = (Response to preferred amplitude - Sum of Responses to the other amplitudes) / (Response to preferred amplitude + Sum of Responses to the other amplitudes)). **m)** Average percentage of shared WR neurons plotted as a function of the distance between stimulus amplitudes. **n)** Support Vector Machine (SVM) multi-class decoding accuracy across 5-frames-sliding-window averages for distinct amplitudes. Gray shaded area indicates accuracy for shuffled data; dotted line is the theoretical chance level. **o)** Time-resolved classification probabilities illustrating misclassification predominantly assigned to adjacent amplitudes. Shaded areas and error bars represent SEM.

**Extended Data Fig. 2.**
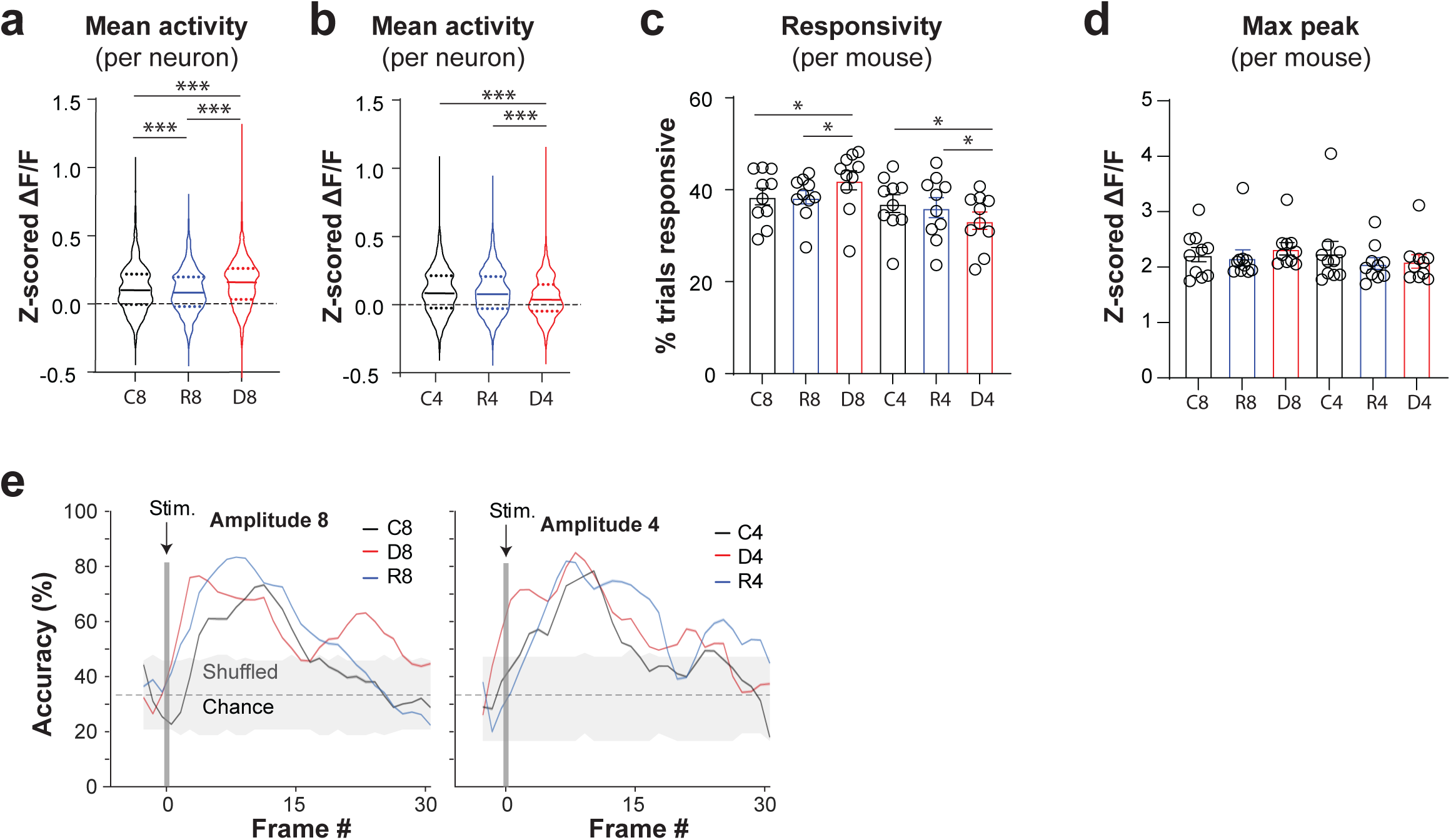
Characterization of amplitude predictability encoding. **a)** Statistical comparison of mean evoked population activity (z-scored ΔF/F) per neuron for Amplitude 8 across three predictability contexts: Control (C8, black), Repetitive (R8, blue), and Deviant (D8, red). **b)**, Same as *A*, but for Amplitude 4. (Linear Mixed-Effects Model, Context p < 0.001, Amplitude p<0.001, Context x Amplitude interaction p<0.001, n= 1,848 neurons from 10 mice). **c)** Statistical comparison per mouse of average responsivity across contexts for Amplitudes 8 (C8, R8, D8) and 4 (C4, R4, D4) (Two-way ANOVA, Context p= 0.695, Amplitude p = 0.002, Context x Amplitude interaction p= 0.003, N= 10 mice). **d)** Statistical comparison per mouse of maximum evoked activity (z-scored ΔF/F) across contexts for Amplitudes 8 (C8, R8, D8) and 4 (C4, R4, D4) (Two-way ANOVA, Context p= 0.065, Amplitude p= 0.046, Context x Amplitude interaction p= 0.284, N= 10 mice). **e)** Support Vector Machine (SVM) multi-class context decoding accuracy across 5-frames-sliding-window averages for Amplitudes 8 (left, C8, R8, D8) and 4 (right, C4, R4, D4). Gray shaded area for empirically derived chance level, dotted line for theoretical chance level. Individual mice data represented in circles. *p<0.05; ***p<0.001. Shaded areas and error bars represent SEM.

**Extended Data Fig. 3.**
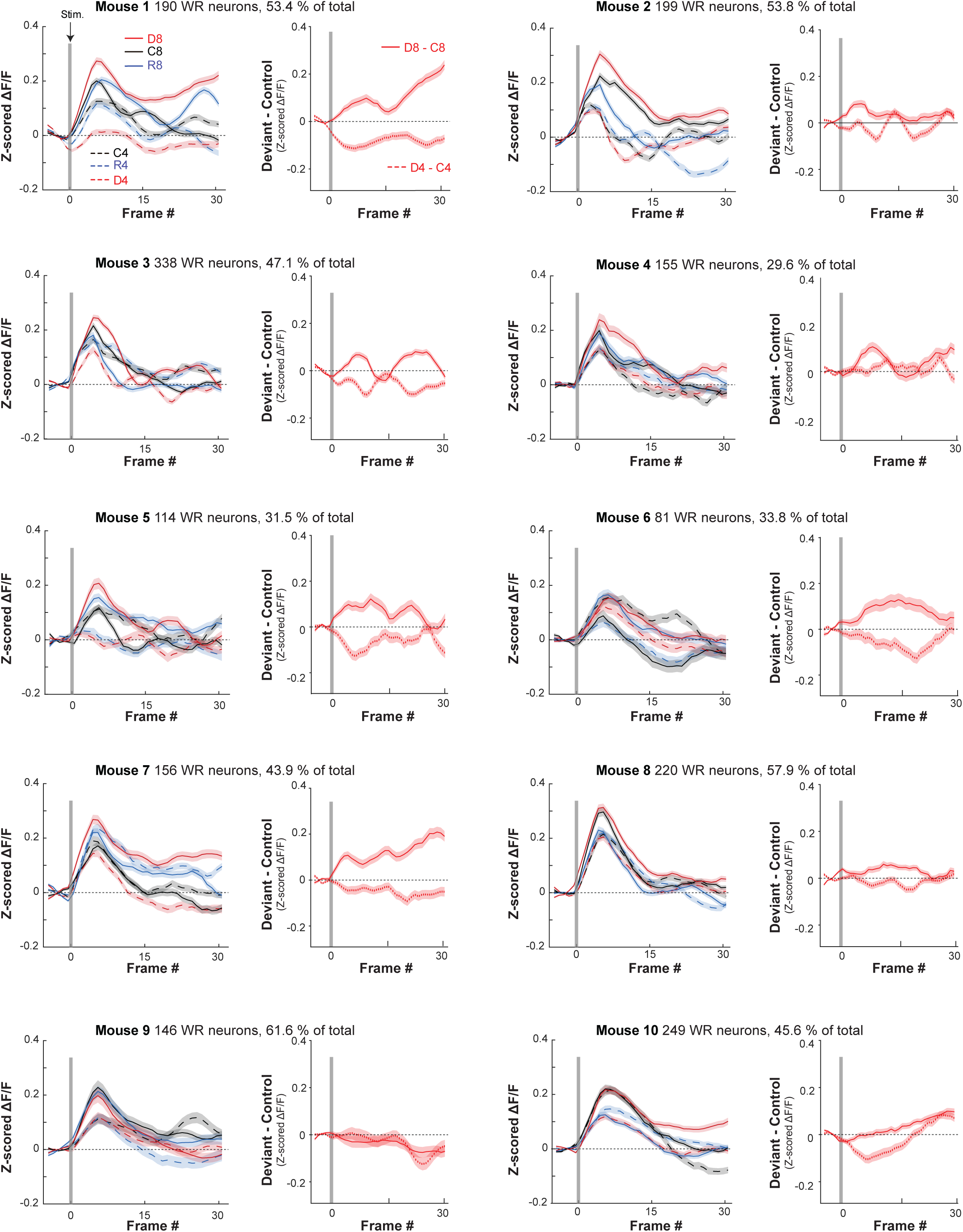
Data for 10 individual mice: *Left*, Mean evoked population activity (z-scored ΔF/F) of all WR neurons per mouse for Amplitude 8 (solid lines) and Amplitude 4 (dashed lines) across three predictability contexts: Control (C, black), Repetitive (R, blue), and Deviant (D, red), contexts. *Right*, subtracted mean evoked population activity (D – C z-scored ΔF/F) isolating the prediction error (PE) modulation (mean evoked response to Deviant minus its corresponding Control) for greater-than-expected (D8 – C8, solid line) and smaller-than-expected violations (D4 – C4, dashed line). Shaded areas represent SEM.

**Extended Data Fig. 4.**
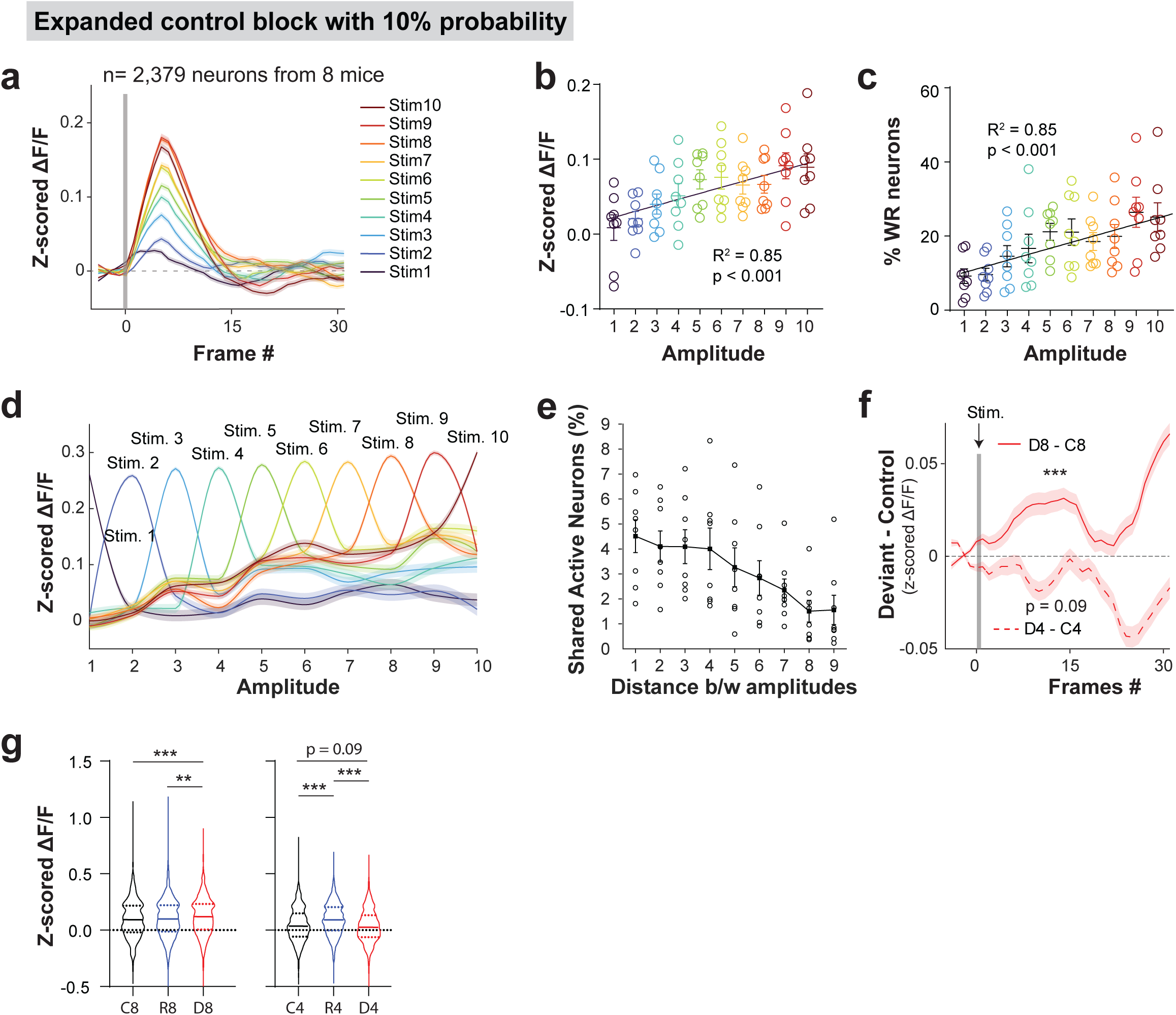
Amplitude predictability encoding in an expanded 10% probability control block. **a)** Mean evoked population activity (z-scored ΔF/F) of WR neurons across the expanded 10-amplitude Control block (n= 2,379 neurons from 8 mice). **b)** Linear correlation between stimulus amplitude and mean evoked population activity per mouse. Individual mice data represented in circles. **c)** Linear correlation between stimulus amplitude and the percentage of recruited WR neurons per mouse. **d)** Tuning curves of the WR neurons to each specific amplitude. **e)** Average percentage of shared WR neurons plotted as a function of the distance between stimulus amplitudes. **f)** Subtracted mean evoked population activity isolating the PE modulation (mean evoked response to Deviant minus its corresponding 10%-occurrence Control) for greater- and smaller-than-expected violations. **g)** Statistical comparison of mean evoked population activity per neuron for Amplitude 8 (left) and Amplitude 4 (right) across three predictability contexts (Linear Mixed-Effects Model, Context p<0.001, Amplitude p<0.001, Context x Amplitude interaction p<0.001, n= 1,878 neurons from 8 mice). *p<0.05; ***p<0.001. Shaded areas and error bars represent SEM.

**Extended Data Fig. 5.**
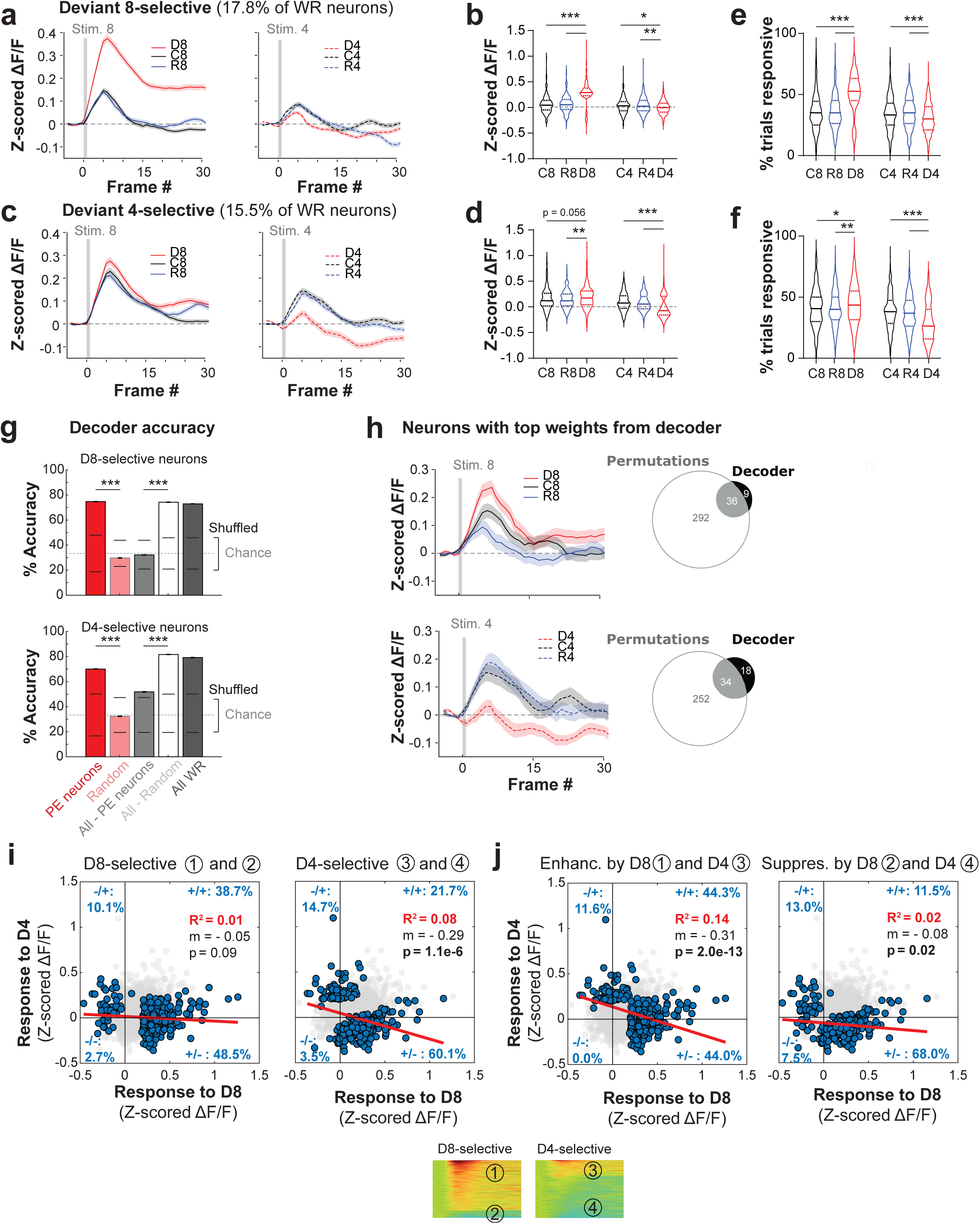
Characterization of pPE and nPE ensembles. **a)** Mean evoked activity (z-scored ΔF/F) of D8-selective neurons (17.8% of WR neurons) in different contexts to Amplitude 8 (left; C8, black; R8, blue; D8, red) and Amplitude 4 (right; C4, R4, D4). **b)** Statistical comparison of D8-selective neurons mean evoked activity across contexts for Amplitude 8 and Amplitude 4 (Linear Mixed-Effects Model, Context p<0.001, Amplitude p<0.001, Context x Amplitude interaction p<0.001, n= 328 neurons from 10 mice). **c)** Same as *a* but for D4-selective neurons (15.5% of WR neurons). **d)** Same as *b* but for D4-selective neurons (Linear Mixed-Effects Model, Context p<0.003, Amplitude p<0.001, Context x Amplitude interaction p<0.001, n= 286 neurons from 10 mice). **e-f)** Statistical comparison of average responsivity per neuron for D8- and D4-selective neurons, respectively, across contexts (Linear Mixed-Effects Model; *E*, Context p<0.001, Amplitude p<0.001, Context x Amplitude interaction p<0.001, n= 328 neurons from 10 mice. *F*, Context p= 0.001, Amplitude p<0.001, Context x Amplitude interaction p<0.001, n= 286 neurons from 10 mice). **g)** Support Vector Machine multi-class context decoding accuracy of mean evoked activity (frames 3-15) across Amplitude 8 (top) or 4 (bottom), for Deviant-selective neurons (red column), an equal number of randomly selected neurons (light red column), the entire WR population minus Deviant-selective neurons (grey column), the entire WR population minus an equal number of randomly selected neurons (light grey column), and the entire WR population (dark grey column). Black lines indicate empirically derived chance level, dotted lines indicate theoretical chance level (t-test, p<0.001, N= 200 iterations for all comparisons). **h)** Mean evoked activity (z-scored ΔF/F) of neurons with the top weights from the decoder trained on the entire population in response to Amplitude 8 (top) and Amplitude 4 (bottom), with accompanying Venn diagrams illustrating the overlap with our selected D-preferring neurons. **i)** Cross-deviant correlation scatter plots for all D8-selective (left) and D4-selective (right) neurons. Percentages indicate the relative distribution of neurons across quadrants. **j)** Cross-deviant correlation scatter plots for neurons enhanced by D8 and by D4 (left), and for neurons suppressed by D8 and by D4 (right). *p<0.05; **p<0.01; ***p<0.001. Shaded areas and error bars represent SEM.

**Extended Data Fig. 6.**
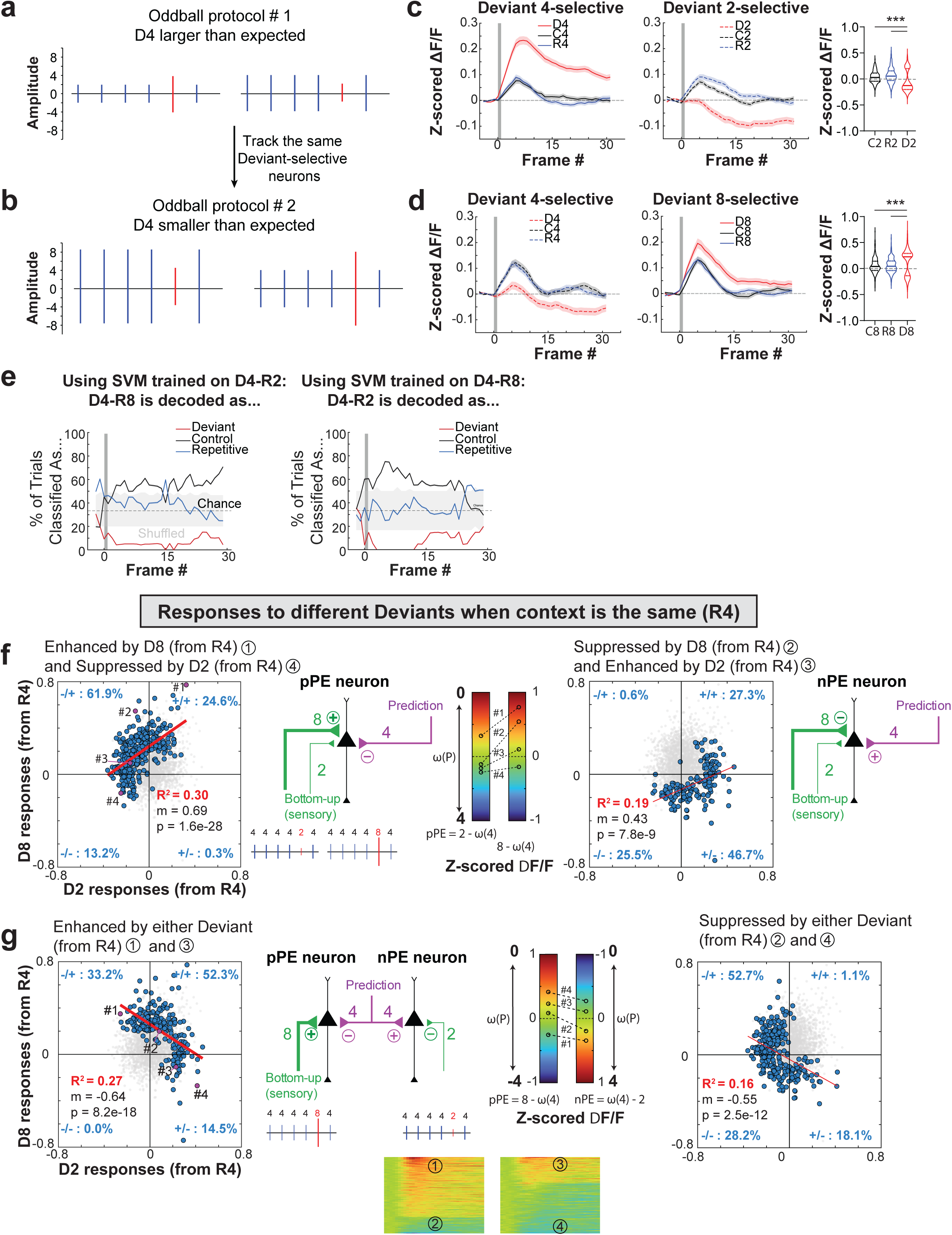
Characterization of pPE and nPE ensembles during fixed sensory and predictive inputs. **a)** Left, Cartoon of the Oddball 1 protocol where D4 is greater than the expected R2 (D4 after R2); Right, flipped oddball (D2 after R4). **b)** Left, Cartoon of the Oddball 2 protocol where the same D4 is now smaller than the expected R8 (D4 after R8); Right, flipped oddball (D8 after R4). **c)** Mean evoked activity (z-scored ΔF/F) of D4-selective neurons chosen after R2 (11.2% of WR neurons) in response to Amplitude 4 (left) and D2-selective neurons chosen after R4 (10.7% of WR neurons) in response to Amplitude 2 (middle), and statistical comparison per neuron (right) (Linear Mixed-Effects Model, Context p<0.001, Amplitude p<0.001, Context x Amplitude interaction p<0.001, n= 286 neurons from 6 mice). **d)** Same as *c* but for D4-selective neurons chosen after R8 (9.2% of WR neurons) and D8-selective neurons chosen after R4 (9.9% of WR neurons) (Linear Mixed-Effects Model, Context p<0.001, Amplitude p<0.001, Context x Amplitude interaction p= 0.001, n= 235 neurons from 6 mice). **e)** Support Vector Machine cross-context decoding probability across 5-frames-sliding-window averages showing decoder generalization across distinct oddball paradigms. Decoder trained on Amplitude 4 during greater-than-expected context (Oddball 1, D4 after R2) and evaluated with D4-evoked neuronal responses during smaller-than-expected context (Oddball 2, D4 after R8) (left). Decoder trained on Amplitude 4 during smaller-than-expected context (Oddball 2, D4 after R8) and evaluated with D4-evoked neuronal responses during greater-than-expected context (Oddball 1, D4 after R2) (right). **f-g)** Cross-deviant correlation scatter plots for the distinct group of neurons indicated. Percentages indicate the relative distribution of neurons across quadrants. Schematic diagrams illustrating the proposed arithmetic framework for pPE and nPE neurons. Heatmap of mean evoked activity from example neurons in panels *f-g*. ***p<0.001. Shaded areas and error bars in *c-d* represent SEM.

**Extended Data Fig. 7.**
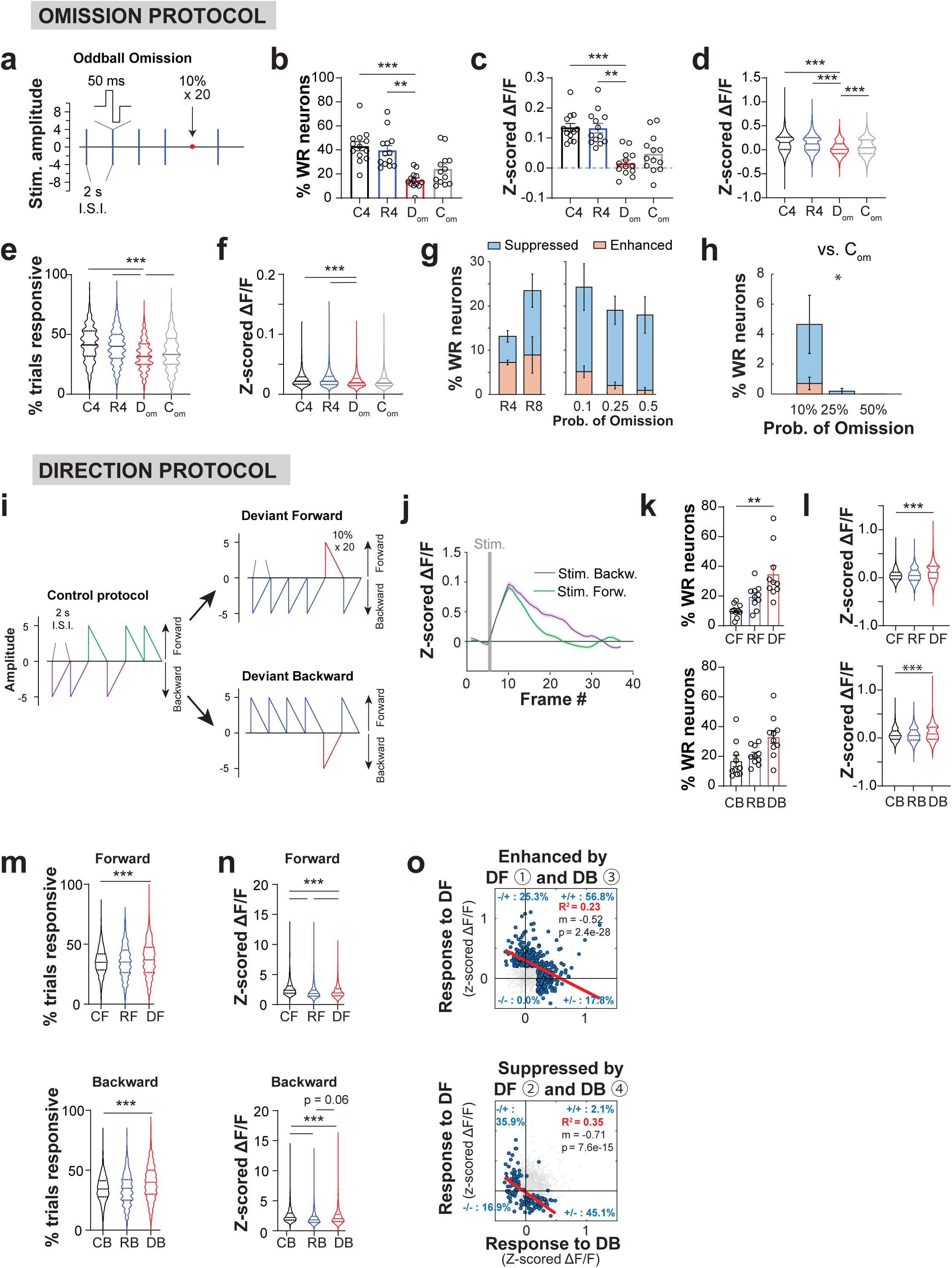
Characterization of sensory omission and direction oddball paradigms. **a)** Cartoon of the Oddball Omission protocol. **b)** Statistical comparison of the percentage of recruited WR neurons per mouse across contexts (C4, R4, C_om_, D_om_) (RM one-way ANOVA, p<0.001, N= 13 mice). **c-d)** Statistical comparison of mean evoked population activity (z-scored ΔF/F) per mouse (*c*, Friedman test, p<0.001, n = 13 mice) and per neuron (*d*, Linear Mixed-Effects Model, Context p<0.001, n = 2,254 neurons from 13 mice) for Control Stimulus (C4, black), Repetitive Stimulus (R4, blue), Control Omission (C_om_, grey), and Deviant Omission (D_om_, red). **e-f)** Statistical comparison per mouse of average responsivity (*e*, Linear Mixed-Effects Model, Context p<0.001, n= 2,254 neurons from 13 mice) and maximum response (*f*, Linear Mixed-Effects Model, Context p<0.001, n= 2,254 neurons from 13 mice). **g)** Statistical comparison of enhanced and suppressed D_om_-selective neurons per mouse as a percentage of WR neurons during the omission of R4 and R8 (left) (Wilcoxon test, all D_om_-selective neurons: p=0.125; enhanced: p= 0.875; suppressed: p= 0.250, N= 4 mice) and across different omission probabilities (*right*) (Friedman test, all D_om_-selective neurons: p=0.653; enhanced: p= 0.278; suppressed: p= 0.431, N = 4 mice). **h)** Statistical comparison of enhanced and suppressed D_om_-selective (C_om_ vs. D_om_) neurons per mouse as a percentage of WR neurons across different omission probabilities (Friedman test, all D_om_-selective neurons: p=0.037; enhanced: p= 0.333; suppressed: p= 0.111, N= 4 mice). **i)** Cartoon of the Oddball Direction protocol. **j)** Mean evoked population activity of all WR neurons during the Control block (n = 1,472 neurons from 10 mice). **k)** Statistical comparison of percentage of recruited WR neurons (Two-way ANOVA, Context p= 0.354, Direction p= 0.006, Context x Direction interaction p= 0.002, n= 10 mice). **l-n)** Statistical comparison across different contexts, per neuron, of mean evoked population activity (*l*, Linear Mixed-Effects Model, Context p<0.001, Direction p= 0.468, Context x Direction interaction p<0.001, n= 1,472 neurons from 10 mice), average responsivity (*m*, Linear Mixed-Effects Model, Context p<0.001, Direction p= 0.762, Context x Direction interaction p < 0.001, n = 1,472 neurons from 10 mice), and maximum evoked activity (*n*, Linear Mixed-Effects Model, Context p<0.001, Direction p= 0.977, Context x Direction interaction p<0.001, n= 1,472 neurons from 10 mice). **o)** Cross-deviant correlation scatter plots for the distinct group of neurons indicated (① enhanced DF group, ② suppressed DF group, ③ enhanced DB group, ④ suppressed DB group). Percentages indicate the relative distribution of neurons across quadrants. Individual mice data represented in circles. **p<0.01; ***p<0.001. Shaded areas and error bars represent SEM.

**Extended Data Fig. 8.**
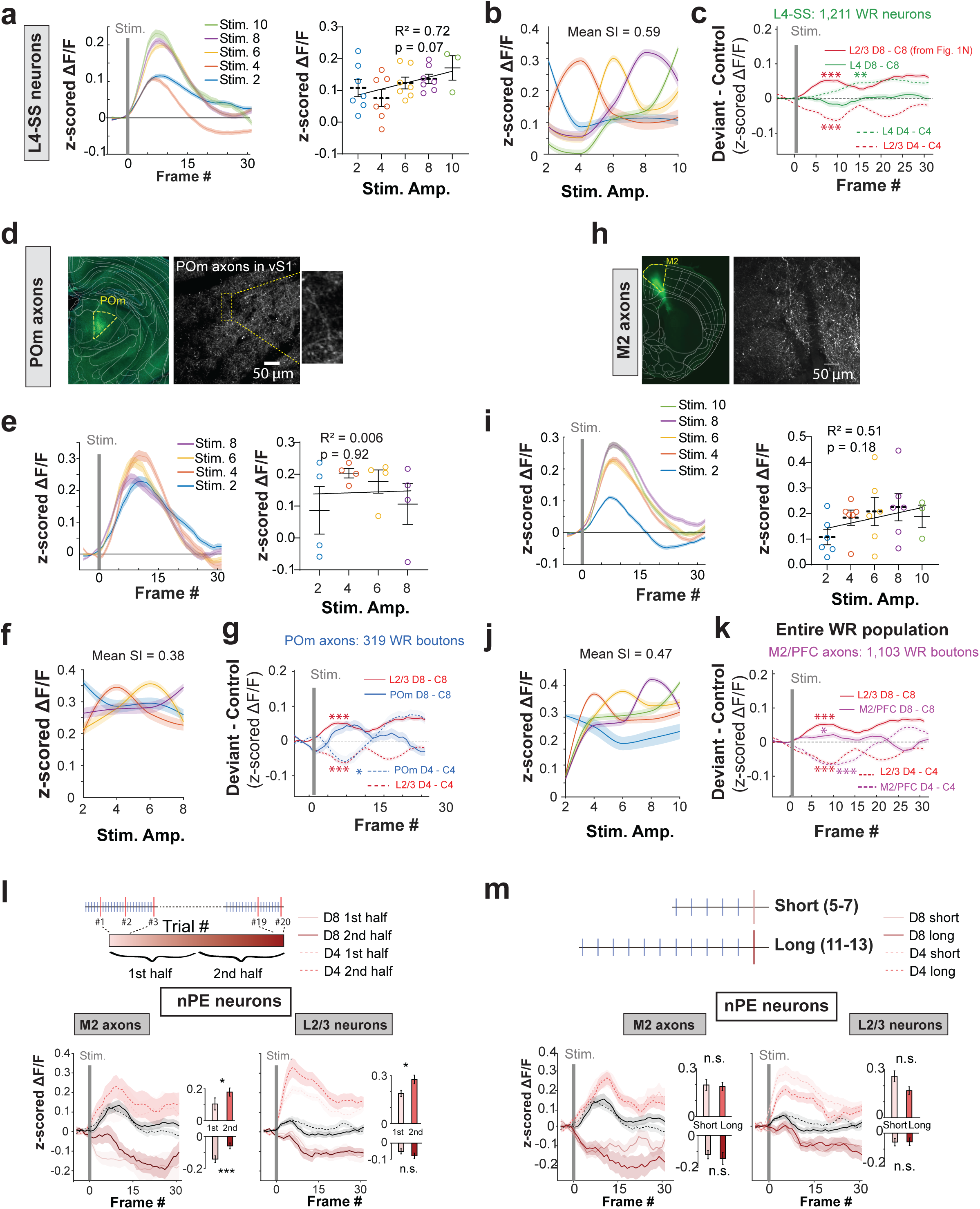
Characterization of prediction error L4 spiny stellate (SS) neurons, POm→S1 and M2→S1 axon boutons and iceberg model. **a)** *Left*, mean evoked population activity (z-scored ΔF/F) of all WR L4 SS neurons relative to all control stimulation amplitudes (n= 974 neurons from 7 mice). *Right*, Linear correlation between stimulus amplitude and mean evoked population activity per mouse (Simple linear regression, R^2^ = 0.718, p = 0.069, N = 7). Individual mice data represented in circles. **b)** Tuning curves of all WR L4 SS neurons to each specific amplitude. **c)** Subtracted mean evoked PE activity (D – C, z-scored ΔF/F) of all WR L4 spiny stellate (SS) neurons to greater- and smaller-than-expected Deviants (Context p<0.001, Amplitude p<0.001, Context x Amplitude interaction p<0.001, n= 1,211 neurons from 7 mice). Equivalent data from L2/3 neurons from Fig. 1n shown for comparison. **d)** Left, histological verification of virus injections targeted to the posteromedial thalamic nucleus (POm). Right, representative in vivo 2-photon calcium imaging field of view showing axons expressing GCaMP8s in vS1 (scale bar = 50 µm) and magnified view of axon boutons (right). **e)** *Left*, mean evoked population activity (z-scored ΔF/F) of all WR POm→S1 axon boutons relative to all control stimulation amplitudes (n= 257 boutons from 4 mice). *Right*, Linear correlation between stimulus amplitude and mean evoked population activity per mouse (Simple linear regression, R^2^ = 0.006, p = 0.923, N = 4). **f)** Tuning curves of all WR POm→S1 axon boutons to each specific amplitude. **g)** Subtracted mean evoked PE activity (D – C) for all WR POm→S1 boutons (Context p=0.020, Amplitude p=0.244, Context x Amplitude interaction p=0.005, n= 319 neurons from 4 mice). **h)** Left, histological verification of virus injections targeted to secondary motor cortex (M2). Right, representative in vivo 2-photon calcium imaging field of view showing axons expressing GCaMP8s in vS1. **i)** *Left*, mean evoked population activity (z-scored ΔF/F) of all WR M2→S1 axon boutons relative to all control stimulation amplitudes (n= 871 boutons from 6 mice). *Right*, Linear correlation between stimulus amplitude and mean evoked population activity per mouse (Simple linear regression, R^2^ = 0.508, p = 0.177, N = 6). **j)** Tuning curves of all WR M2→S1 axon boutons to each specific amplitude. **k)** Subtracted mean evoked PE activity (D – C) for all WR M2→S1 boutons (Context p<0.001, Amplitude p<0.001, Context x Amplitude interaction p<0.001, n= 1,103 neurons from 6 mice). **l)** Trial-by-trial mean evoked activity (z-scored ΔF/F) for the first half Deviant trials and statistical comparison (insets) for D8 (dark red) and D4 (light red) in putative nPE M2-to-S1 axons (*left*) (Amplitude 8: Paired t-test, p<0.001; Amplitude 4: Paired t-test, p= 0.049, n= 72 from 6 mice) and L2/3 pyramidal neurons (*right*) (Amplitude 8: Paired t-test, p= 0.165; Amplitude 4: Paired t-test, p= 0.013, n= 112 from 10 mice). **m)** Short-term experience-dependent mean evoked activity (z-scored ΔF/F) and statistical comparison (insets) for short (5–7, light red) and long (11–13, dark red) repetitive sequences preceding a deviant stimulus in putative nPE M2-to-S1 axons (*left*) (Amplitude 8: Paired t-test, p= 0.498; Amplitude 4: Paired t-test, p= 0.814, n= 72 from 6 mice) and L2/3 pyramidal neurons (*right*) (Amplitude 8: Wilcoxon test, p= 0.525; Amplitude 4: Wilcoxon test, p = 0.093, n= 112 from 10 mice). *p<0.05; **p<0.01; ***p<0.001. Shaded areas and error bars represent SEM

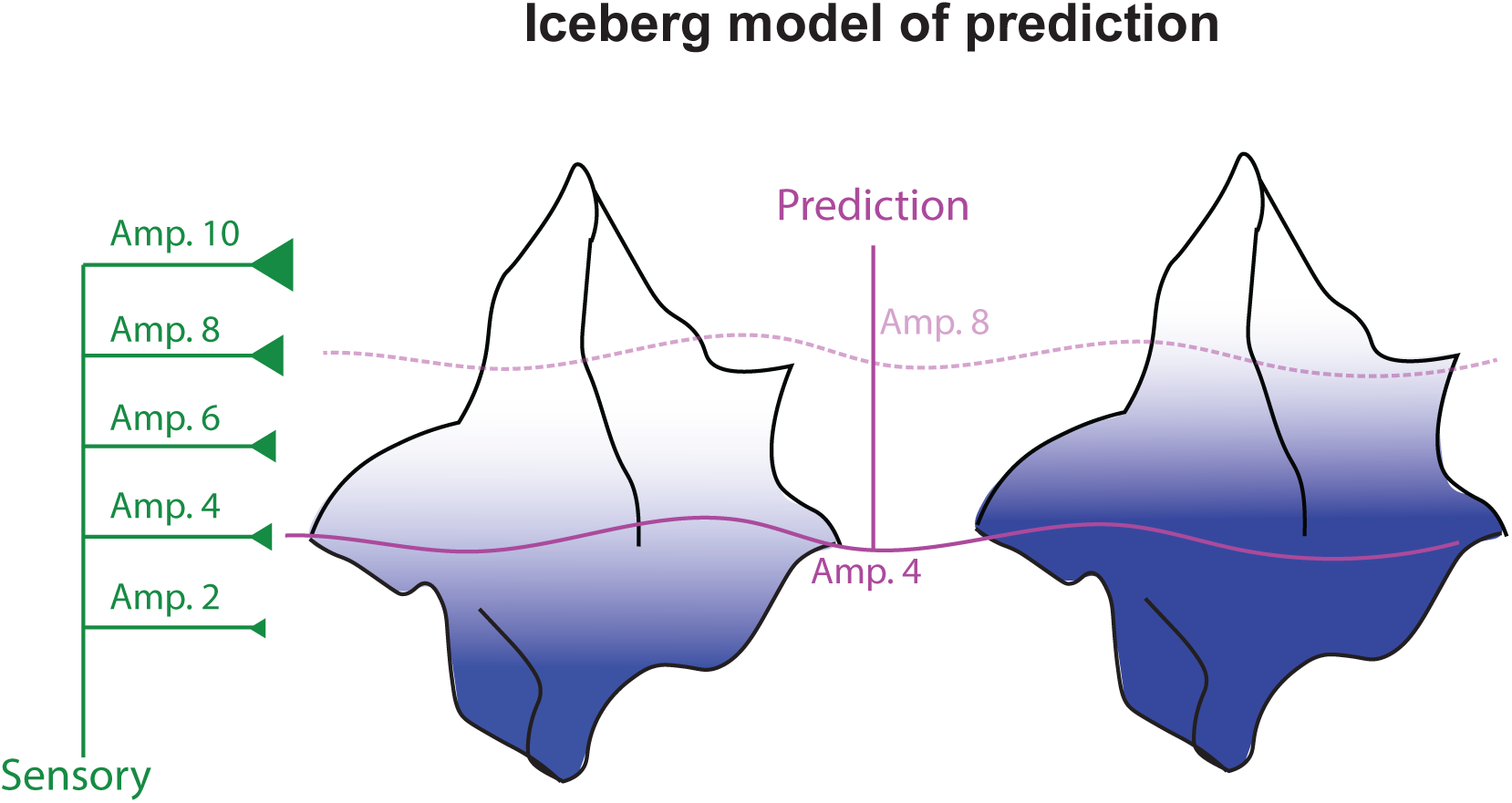

